# Meningioma transcriptomic landscape demonstrates novel subtypes with regional associated biology and patient outcome

**DOI:** 10.1101/2024.02.23.581766

**Authors:** H. Nayanga Thirimanne, Damian Almiron Bonnin, Nicholas Nuechterlein, Sonali Arora, Matt Jensen, Carolina A. Parada, Chengxiang Qiu, Frank Szulzewsky, Collin W English, William C Chen, Philipp Sievers, Farshad Nassiri, Justin Z. Wang, Tiemo J Klisch, Kenneth D Aldape, Akash J Patel, Patrick J Cimino, Gelareh Zadeh, Felix Sahm, David R. Raleigh, Jay Shendure, Manuel Ferreira, Eric C Holland

**Affiliations:** Human Biology Division, Fred Hutchinson Cancer center, Seattle, WA, USA; Department of Pathology, University of California San Francisco, USA; Neuropathology Unit, Surgical Neurology Branch, National Institute of Neurological Disorders and Stroke, National Institutes of Health, Bethesda, MD, USA; Seattle Translational Tumor Research Center, Fred Hutchinson Cancer Center, Seattle, WA, USA; Department of Neurological Surgery, University of Washington Medical Center, Seattle, WA, USA; Department of Genome Sciences, University of Washington, Seattle, WA, USA; Department of Neurosurgery, Baylor College of Medicine, Houston, TX, USA; Departments of Radiation Oncology, Neurological Surgery, and Pathology, University of California San Francisco, USA; Department of Neuropathology, Institute of Pathology, University Hospital Heidelberg, Heidelberg, Germany; Clinical Cooperation Unit Neuropathology, German Consortium for Translational Cancer Research (DKTK), German Cancer Research Center (DKFZ), Heidelberg, Germany; Department of Surgery, Division of Neurosurgery, University of Toronto, Toronto, ON, Canada; Laboratory of Pathology, Center for Cancer Research, National Cancer Institute, Bethesda, MD, USA

**Keywords:** Meningioma, brain tumor, UMAP, bulk RNA-Seq, Oncoscape, patient prognosis prediction, meningioma subtypes, recurrent

## Abstract

Meningiomas, the most common intracranial tumor, though mostly benign can be recurrent and fatal. WHO grading does not always identify high risk meningioma and better characterizations of their aggressive biology is needed. To approach this problem, we combined 13 bulk RNA-Seq datasets to create a dimension-reduced reference landscape of 1298 meningiomas. Clinical and genomic metadata effectively correlated with landscape regions which led to the identification of meningioma subtypes with specific biological signatures. Time to recurrence also correlated with the map location. Further, we developed an algorithm that maps new patients onto this landscape where nearest neighbors predict outcome. This study highlights the utility of combining bulk transcriptomic datasets to visualize the complexity of tumor populations. Further, we provide an interactive tool for understanding the disease and predicting patient outcome. This resource is accessible via the online tool Oncoscape, where the scientific community can explore the meningioma landscape.

## Introduction

Meningiomas are the most common primary intracranial tumors in humans and usually benign. However, some are malignant, rapidly recur after multimodal treatment with surgery and radiotherapy, and can ultimately be fatal^1^. The histologic grading of the 2021 World Health Organization (WHO) classification of tumors of the central nervous system identifies many of these malignant tumors, but some tumors identified as grades 1 or 2 are equally aggressive^2^. Improved risk classification systems for these tumors are needed, and several grading systems based on DNA methylation, copy number, or expression signatures have been proposed^1,3–5^.

Clues to the underlying biology of these tumors come from neurofibromatosis type 2 with germline loss of one copy of *NF2*, resulting in the formation of multiple meningiomas^6^. Consistent with this observation, DNA molecular analysis shows that the most common alteration of spontaneous meningiomas is loss of chromosome 22 harboring the *NF2* gene^7,8^. The majority of rapidly recurrent meningiomas are among those that show functional loss of *NF2*^9,10^. Additional recurrent genetic alterations in *NF2*-wildtype meningiomas include mutations in genes *TRAF7*, *KLF4*, *AKT1* and *SMO*, and are often restricted to benign meningiomas^11,12^.

*NF2*, encodes the protein Merlin which is a tumor suppressor that regulates YAP1 via the Hippo signaling pathway^7^. Upon contact inhibition, the Hippo pathway phosphorylates YAP1, resulting in the inhibition of YAP1 activity^13^. In the absence of Merlin, YAP1 remains active and translocate into the nucleus, binding the TEAD transcription factors, and activating cell proliferation. In addition to chromosome 22 loss, the remaining *NF2* allele is frequently inactivated due to either inactivating point mutations in the *NF2* sequence or gene fusions involving the *NF2* gene. An alternative route of YAP1 de-regulations can occur due to gene fusions involving the YAP1 gene, resulting in constitutively active YAP1 that is insensitive to Hippo pathway inactivation^14^. Mouse modeling experiments have shown that the expression of either constitutively active YAP1 or YAP1 gene fusions found in human meningiomas induce similar tumors in mice, functionally implicating de-regulated YAP activity in the pathobiology of meningiomas^15^.

Currently available therapeutic options for patients with aggressive meningiomas are limited to radiation and multiple surgeries, therefore a better understanding of the underlying biology of aggressive meningiomas is needed. It is likely that rapid recurrence and aggressive behavior of some meningiomas reflects the underlying biology of these tumors, which is in turn largely reflected by its overall gene expression pattern. In the hope of understanding this aggressive subset of meningiomas and being able to predict which meningiomas will fall into that category, we performed RNA sequencing of 279 meningiomas from all grades. We then combined our data with multiple publicly available meningioma RNA-Seq datasets to generate one of the largest clinically annotated meningioma full RNA-Seq datasets available to define the biology of the various meningioma subgroups to create a reference landscape using Uniform Manifold Approximation and Projection (UMAP) of 1298 tumors with associated metadata.

This reference map exhibits multiple clusters of tumors each represented by multiple datasets. We find that there are multiple RNA-Seq based meningioma subtypes, some of which are associated with distinct time to recurrence, that are distinguished from each other by gene expression similarities to developmental cell types and biological pathways. We observed several subtypes with particularly poor outcomes; the most common aggressive tumors of these showed high proliferation rates and RNA expression resembling muscle development. We also sought to develop a method to map new patients onto our UMAP landscape and predict tumor behavior and patient outcome based on the nearest neighboring tumors in the map. Oncoscape, an open-source online tool via which this reference map is accessible not only provides a great visualization platform for the data shared in this article, but it also allows interactive and analytical exploration of tumors along with various associated metadata. (Oncoscape is accessible on the Chrome search engine via the link: https://oncoscape.sttrcancer.org/#project_meningiomaumap91. The main figure panels can be directly accessed using the dropdown menu “Figures from the paper” on upper-right corner of the website.) We believe that this reference map with demographic and clinical data provides insight to the behavior of the multiple meningioma subtypes, and tools to map new patients on to it will be beneficial in clinical applications to determine patient outcome and therapeutic strategies.

## Results

### Constructing the meningioma reference UMAP

We obtained 12 publicly available bulk RNA-Seq meningioma datasets from nine institutions and 5 countries in North America, Europe, and Asia and combined them with 279 sequenced meningiomas from the University of Washington to create a set of 1298 meningiomas (Table S1)^3,4,14,16–25^. Raw sequencing data were collected from each dataset and aligned to human genome hg38 using the same pipeline (Figure 1A). To remove batch effects from different datasets, we used R function CombatSeq from the R package “sva”. Gene expression values from combined datasets were normalized and converted to units of transcripts per million (TPM)^26^. We applied Uniform Manifold Approximation and Projection (UMAP), a dimensionality reduction method, on batch-corrected, normalized transcript counts to create a reference UMAP (Figure 1A-B, Figure S1A). This landscape is made up of multiple clusters of different sizes, all of which are composed of a mix of the 13 datasets with the exception of the HKU/UCSF dataset (GSE212666) for which a minor subset of patients forms two, small unique clusters (11% of HKU/UCSF dataset) (Figure 1B). In addition to UMAP, we explored other dimension reduction techniques (Principal Component Analysis (PCA) and t-distributed Stochastic Neighbor Embedding (tSNE)) and found that UMAP better distinguished clusters that showed differences in clinical and genomic features (Figures S1B, S1C)^27^. The collection of tumor samples included fresh frozen tissue as well as Formalin-Fixed Paraffin-Embedded tissue. The FFPE samples distributed evenly across the landscape (Figure S1D). The UMAP landscape facilitates 2D- or 3D-visualization and is available for interactive analysis and visualization via the open source, interactive online tool Oncoscape^28^ (see Figure 1 in Oncoscape).

**Figure 1.**
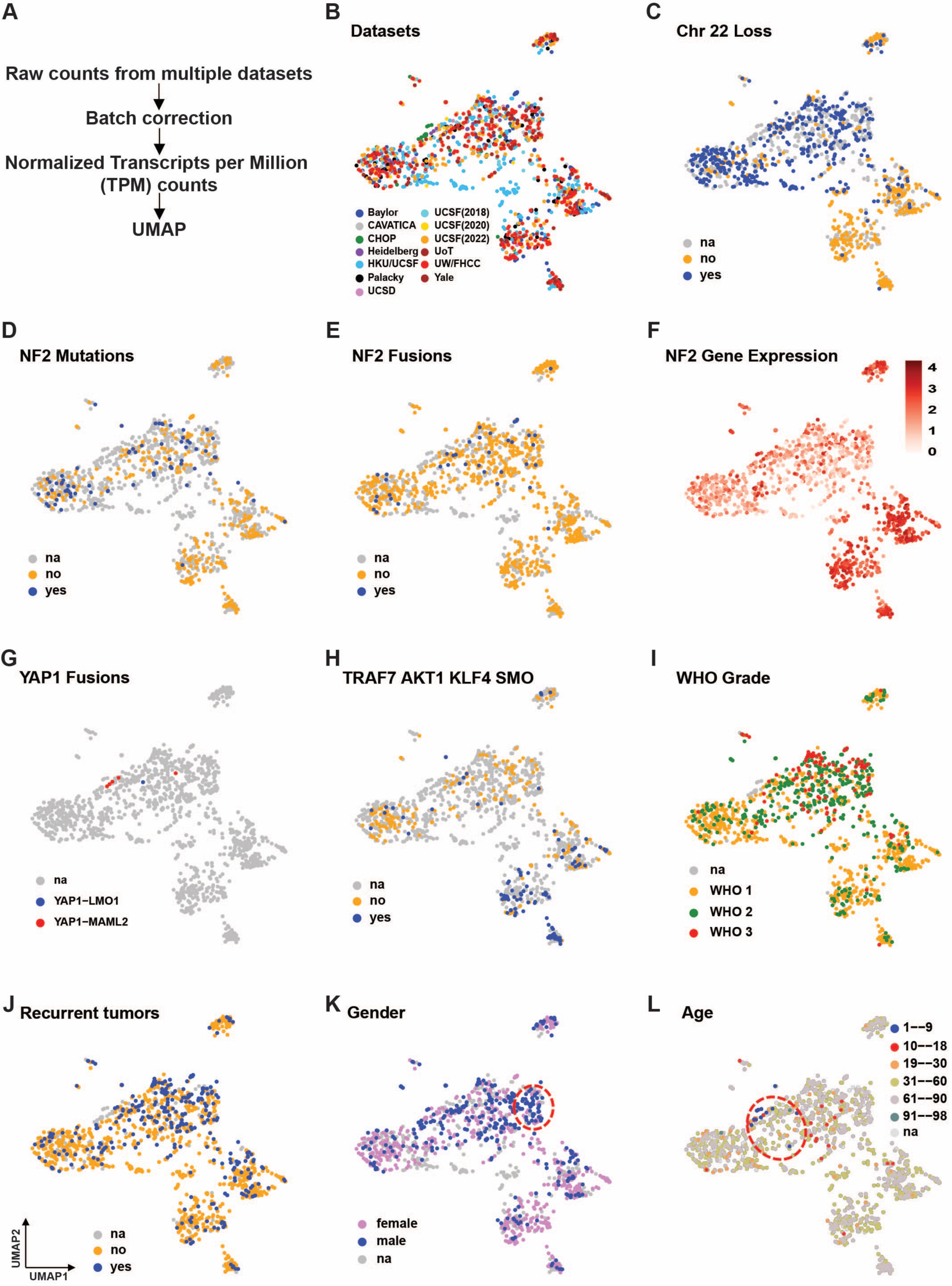
Generating the meningioma reference UMAP and coloring by clinical and genomic metadata. (A) Overview of the method. (B) UMAP colored in by datasets used in this study (C) Tumors with (in blue) and without (in yellow) loss of chromosome 22. Tumors with (in blue) and without (in yellow) (D) *NF2* mutations and (E) *NF2* gene fusions. (F) Expression of *NF2.* (G) Tumors with YAP1 gene fusions (H) Tumors with (in blue) and without (in yellow) mutations in *TRAF7 / KLF4 / AKT1 / SMO* (I) WHO grade of tumors; grade 1 (yellow), 2 (green) and 3 (red) (J) Recurred (in blue) and primary (in yellow) tumors (K) Patients’ gender (female in pink and male in blue). Region 1 marked by the red dashed line. (L) Patients’ age at the time of sample acquisition. Region 2 marked by the red dashed line. (na = not available). (see Figure 1 in Oncoscape) See also Figure S1 and Table S1.

### Known genetic aberrations are regionally distributed across the UMAP

We colored in the UMAP by DNA sequencing-based mutational metadata associated with a subset of the tumors. For any coloring scheme we colored in only known values; tumors with no known values for a given data field were left gray. More than half of meningiomas (73% of tumors for which *NF2* status is available) exhibit functional loss of *NF2*, which is achieved via either the loss of chromosome 22, point mutations, or gene fusions. We colored the map for tumors with known chromosome 22 loss (from collected metadata), which clearly highlighted a large subregion of the map (Figure 1C). Point mutations and gene fusions leading to inactivation of *NF2* also cluster with the tumors with chr22 loss (Figure 1D, 1E). Coloring the UMAP for all three mechanisms of *NF2* inactivation (chr22 loss, mutations, fusions) demonstrated a near complete loss of *NF2* across this region of the map, which is characterized by overall downregulated expression of *NF2* (Figure 1F, Figure S1E). *NF2* wild-type YAP1 fusion-positive meningiomas also mapped on to the same region, indicating that they resemble *NF2* mutant meningiomas on a gene expression level (Figure 1G)^15,29^. Other recurrent non-*NF2* mutations including *TRAF7* and *SMO* were found distributed across the *NF2*-wildtype region of the map, while *KLF4* and *AKT1* additionally showed high regionality for recurrent mutations (Figure 1H, Figure S1F). The regionality of these genetic alterations is consistent with the known unique biology for the meningiomas. For example, meningiomas that harbor both *TRAF7* and *KLF4* mutations were predominantly regionalized to one cluster (Figure S1G)^30^. *TRAF7*-mediated cell transformation is enhanced by loss of *KLF4* in a subset of meningiomas^31^.

### Aggressive tumors are regionally concentrated

Most of the samples in our reference dataset have a WHO grade associated with them, and coloring in the UMAP by that grade shows nonrandom distribution. A subset of the region characterized by *NF2* loss had an increased concentration of WHO 2 and 3 tumors relative to the rest of the map (Figure 1I). However, the region with the highest concentration of WHO 2 and 3 tumors still contained tumors of all three grades. For some of the tumors, there was data indicating whether the tumor sample was a first resection or whether it was a recurrent tumor. Coloring in these data on the map showed that tumors known to be recurrent at the time of resection are generally also concentrated in the same region as tumors with a higher grade (Figure 1J). For some of the patients, there were data regarding the time between the surgery generating the sample and the next or previous resection. Coloring in the map with these data also showed that a short time to recurrence was enriched in that same area as was higher average grade and increased likelihood of being a recurrent tumor (Figure S1H).

### Regional age and gender distribution

Nearly all the samples had records of age and gender. Consistent with what is known, the majority of the map comprised of older patients that were predominantly female (median age of 58 years and 66% female) (Figure 1K, 1L)^32,33^. By contrast, there were two regions of the UMAP that varied from this general rule. One region within the most aggressive tumors were largely male (61% of male vs 31% male in rest of the UMAP), and a second adjacent region comprised of a higher percent of younger patients (< 30 years) than the general populations of meningioma patients (22% vs 5% in rest of the UMAP) (Figure S1I, S1J, S1K).

### Various grading systems are consistent with the regional patterns across the UMAP

Because the WHO grading system does not identify all the meningiomas with aggressive behavior, several recent alternative grading systems have been proposed that use methylation patterns, copy number alterations, and gene expression to place patients into specific groups associated with a time to recurrence^3,4,21,34,35^. We colored in this UMAP by metadata of these grading systems, and all of them correlate with UMAP subregions (Figure 2A-2C, Figure S2A) (see Figure 2 in Oncoscape). Additionally, the expression of the 34 genes presented by Raleigh lab as a signature to predict meningioma outcome was analyzed in correlation to our UMAP^36^. Upregulated and downregulated genes in the most aggressive meningiomas were divided into two gene sets, and the whole dataset was subjected to Gene Set Variation Analysis (GSVA) using the two gene sets separately. The two sets of GSVA scores and then a ratio of them were used to color in the UMAP and they highlighted the most aggressive region of our map in line with the UCSF findings (Figures 2D,2E,2F). Furthermore, we generated a UMAP using only those 34 genes (Figure S2B) and found that basic regionalization of aggressive and recurrent tumors was in line with the UMAP generated using all protein coding genes, however meningioma subtypes were better segregated in the latter UMAP. The one classification system for meningiomas that did not correlate with map location was the Simpson grading scale, which is a measure of tumor resection completeness (Figure S2C)^37^. Lack of regional correlation for the Simpson grading scale suggests that the ability to completely resect a tumor is not determined by expression pattern and underlying biology of the tumor.

**Figure 2.**
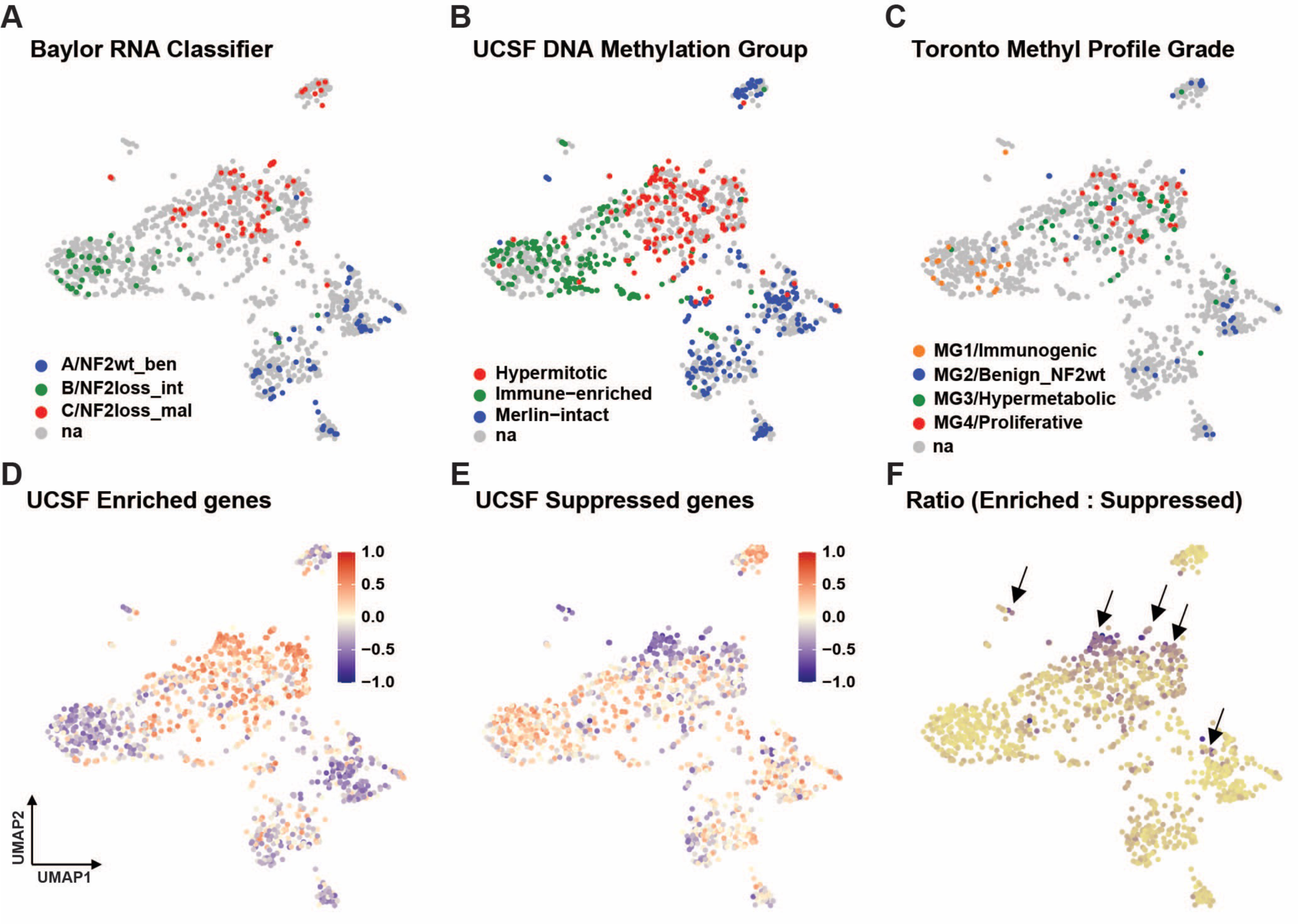
Various grading systems show regional patterns across the UMAP. (A) UMAP colored in by the Baylor RNA classification of a subset of tumors: “A/NF2wt_ben” = NF2 wild-type benign (blue), “B/NF2loss_int” = NF2 lost intermediate (green), “C/NF2loss_mal” = NF2 lost malignant (red). (B) UMAP colored in by UCSF DNA methylation-based classification of a subset of tumors: “Hypermitotic” in red, “Immune-enriched” in green, “Merlin-intact” in blue. (C) UMAP colored in by the Toronto methylation profile of a subset of tumors: “MG1/Immunogenic” in orange, “MG2/Benign_NF2wt” (Benign NF2 wild type) in blue, “MG3/Hypermetabolic” in green, “MG4/Proliferative” in red. (D) UMAP colored in by GSVA scores calculated using UCSF gene set upregulated in most aggressive meningioma. A score closer to 1 suggests upregulation of the respective gene set while a score closer to −1 suggests downregulation of the respective gene set. (E) UMAP colored in by GSVA scores calculated using UCSF gene set downregulated in most aggressive meningioma. A score closer to 1 suggests upregulation of the respective gene set while a score closer to −1 suggests downregulation of the respective gene set. (F) UMAP colored in by the ratio of GSVA scores from upregulated gene set and GSVA scores from downregulated gene set. Black arrows indicate the regions with the most aggressive tumors marked by the ratio of GSVA scores. (see Figure 2 in Oncoscape) See also Figure S2.

### Meningioma subtypes with distinct time to recurrence

Genomic and clinical metadata integrated into the UMAP revealed regionalization, suggesting potential meningioma subtypes. In order to define map regions with statistical confidence, we employed three methods: DBSCAN, k-means, and hierarchical clustering (Figure 3A, Figure S3A, Figure S3B). There was overall overlap among clusters identified by the three methods, however discrepancies existed due to variations in computation methods. The addition of more samples would likely enhance the robustness of the clustering results. We ultimately chose DBSCAN due to its ability to delineate regional distinct clusters that corresponded well with metadata, identifying nine general clusters labeled A through I (Figure 3A) (see Figure 3 in Oncoscape). Notably, both k-means and hierarchical clustering outputs corroborated the UMAP-based intra-cluster subdivisions, as depicted below.

**Figure 3.**
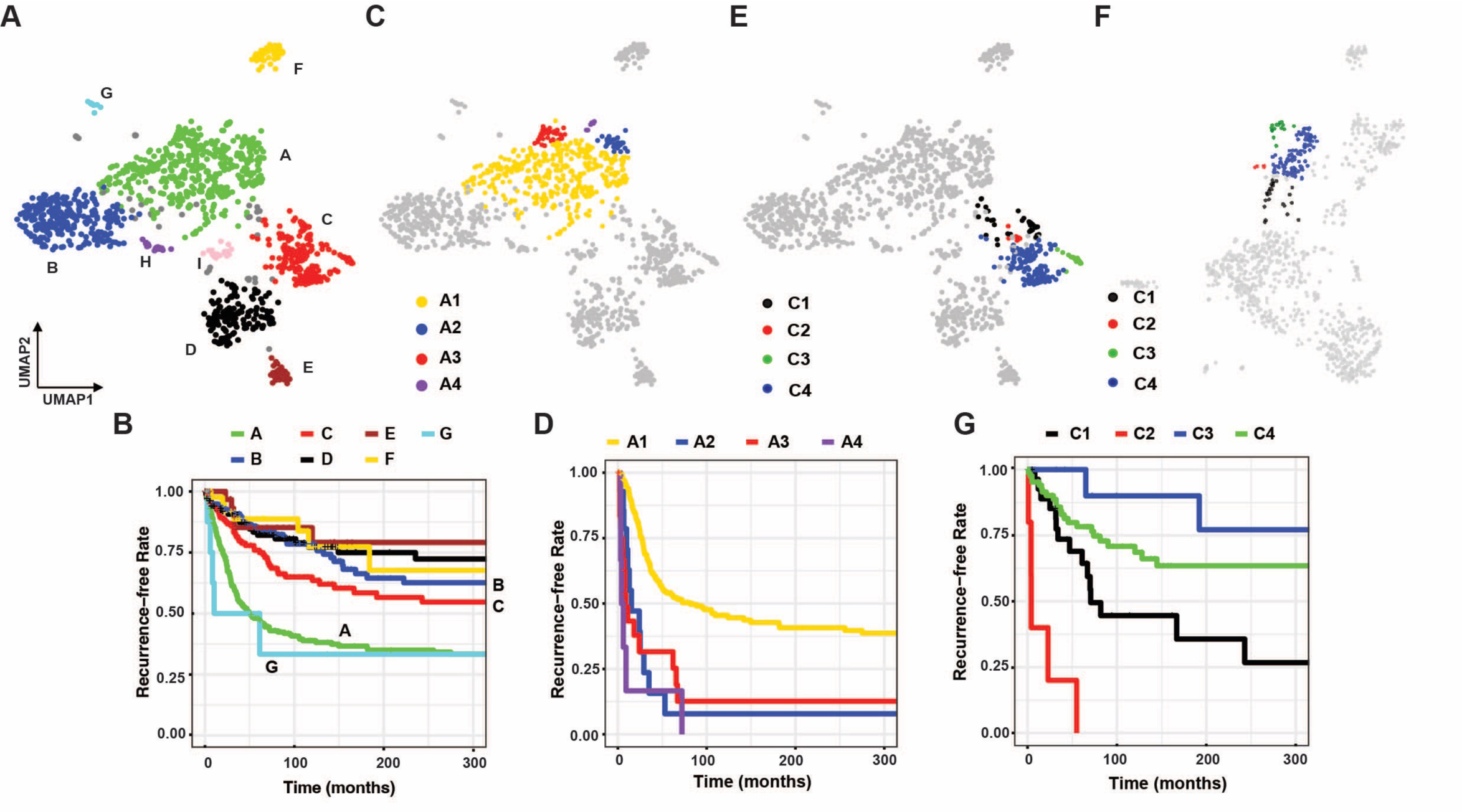
Meningioma subtypes with distinct time to recurrence. (A) Nine major clusters identified by DBSCAN that denotes meningioma subtypes (A to I). Unclustered samples (n=48) are shown in grey. (B) Kaplan Meier plots for the nine clusters based on time to recurrence (AvsB and AvsC p.value <0.0001; CvsD and BvsC p.value < 0.05) (C) Subclusters of cluster A (A1, A2, A3 and A4) (D) Kaplan Meier plots showing recurrence-free rates of cluster A subclusters (A1vsA3: p.value < 0.0001; A1vsA2: p.value < 0.0001; and A2vsA3: p.value = 0.9, A3vsA4: p.value = 0.39) (E) Subclusters of cluster C (C1, C2, C3 and C4) in 2D (F) snapshot of 3D view of cluster C in Oncoscape (G) Kaplan Meier plots showing recurrence-free rates of cluster C subclusters (C1vsC2: p.value < 0.0001, C3vsC4: p.value = 0.26, C1vsC3 p.value = 0.01). (see Figure 3 in Oncoscape) See also Figure S3.

The region of the UMAP with functional loss of *NF2* (Figure S1E) comprised of clusters A and B. Cluster A contained the highest density of aggressive tumors, and cluster B represented the remainder of the *NF2* loss region of the UMAP with relatively benign tumors. Clusters C and D were comprised of mostly *NF2* wild-type tumors. The comparison of Kaplan Meier plots of time to recurrence for these main clusters identified cluster A and G as the clusters with the shortest time to recurrence (Figure 3B). Patients in cluster C also perform significantly worse than those in clusters B, D, E and F, all of which were similar. Clusters H and I comprised of only patients from the Hong Kong dataset.

### Intra-cluster time to recurrence analysis suggests further subdivision of the map

The regional differences of time to recurrence within a cluster was seen for many of the clusters. The most striking was in clusters A and C. We identified four subclusters within cluster A based on the regionalization of the most aggressive tumors and most importantly differences in patient outcome (Figure 3C, 3D). A3 harbored the largest population of patients with poor outcomes. Subcluster A4, although only contains 8 tumors, is derived from 2 datasets, and demonstrated the worse clinical outcomes. Subclusters A2, A3 and A4 were also highlighted as the most aggressive areas by the UCSF gene signature (Figure 2F). Cluster C can be similarly divided into 4 subclusters with significant differences in outcome and these subclusters are prominent when UMAP is visualized in 3 dimensions (Figures 3E-3G). Subcluster C2, like A4, is a small subcluster of 8 tumors derived from 6 different datasets having a uniform short time to recurrence. Several of the other clusters can also be subdivided into regions with significant differences in outcome, however, these clusters represent less aggressive tumor types with few recurrences in general and long times to recurrence (Figure S3C, S3D).

### Gene fusion calling from RNA-Seq data shows high prevalence in aggressive regions

We identified gene fusions from the RNA-Seq data using Arriba^38^. At high confidence we were able to identify 171 gene fusions that have at least one coding gene partner and that recur at least twice within the dataset (Table S4A). The regions of the map with highly aggressive tumors (cluster A and parts of cluster C) showed significantly high fusion burden (Figure 4A) (see Figure 4 in Oncoscape). Some of the tumors harbored multiple fusions and some highly recurrent protein-coding gene fusions were found regionalized on the map (Figure 4B, 4C). For example, TRPM3—TRPM3 fusions enriched within cluster A and some parts of cluster C and F, the most aggressive parts of each cluster. In another example, PARD6B—BCAS4 fusions were enriched in cluster A and most concentrated within the subcluster A3. We identified more *NF2* fusions in addition to what was known from collected metadata, and they were predominant in clusters A and B (Figure S4A). It is worth noting that YAP1 fusions are mostly in pediatric patients (Figure 1G, 1L). YAP1-MAML2 is identified as a causal oncogenic driver in pediatric *NF2* wild-type meningiomas^14^. Furthermore, YAP1-MAML2, which leads to constitutive activation of YAP1, has been shown to be sufficient to induce meningiomas in mice^15^. The pediatric tumors achieve YAP1 activation by different mechanisms with fewer losses of chromosome 22 and more gene fusions such as those that activate YAP1.

**Figure 4.**
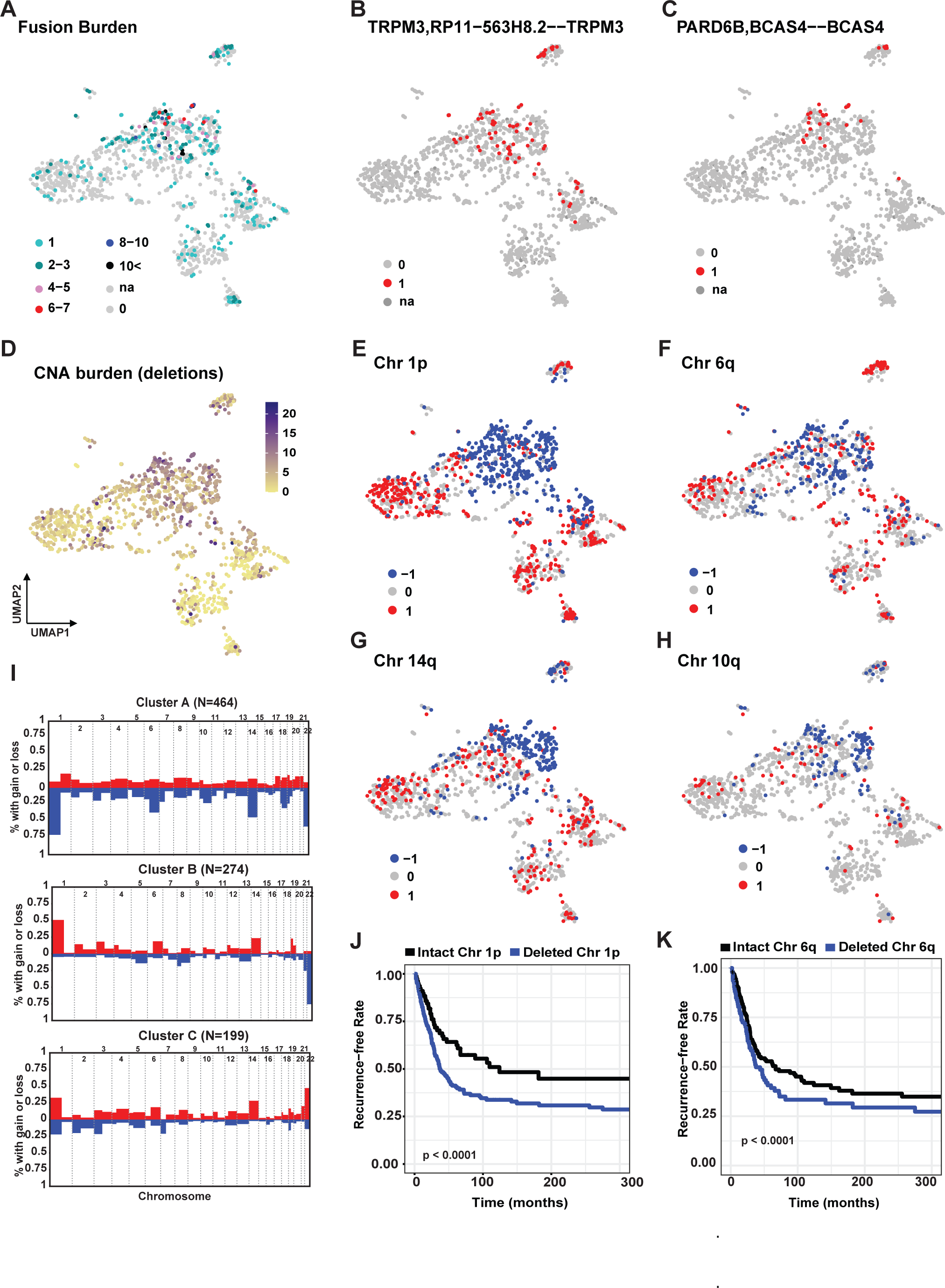
Regionally enriched gene fusions and copy number alterations. (A) Fusion burden in each tumor derived from high confidence gene fusions called using bulk RNA-Seq. (B, C) Examples for regionalized fusions. (D) Burden of Copy Number Alterations (CNA) in each tumor (loss of chromosome arms). (E) Loss (−1), gain (1) or intact (0) status of chromosome 1p in each tumor (F) Loss (−1), gain (1) or intact (0) status of chromosome 6q in each tumor (G) Loss (−1), gain (1) or intact (0) status of chromosome 14q in each tumor (H) Loss (−1), gain (1) or intact (0) status of chromosome 10q in each tumor (I) Manhattan plots showing losses (blue) and gains (red) of each chromosome arm in clusters A, B and (J) Kaplan Meier plot showing the recurrence-free rate of patients in Cluster A with intact and deleted chr 1p and (K) chr 6q. (see Figure 4 in Oncoscape) See also Figure S4, Table S4A and S4B.

### Regionally enriched chromosome arm-level copy number alterations

We estimated arm length gains and losses of chromosomes using CaSpER on RNA-Seq data (Table S4B)^39^. These were validated with the known copy number alteration (CNA) data on 304 samples where DNA sequencing was available for chromosome 22 status, and 90% of 22q loss identified by CaSpER were confirmed by available metadata. Additionally, we show that chromosome 22 losses called by CaSpER correlate well with the expression of *NF2* gene which is harbored on chromosome 22 (Figure S4B). The CNA burden was highest in cluster A where the most aggressive tumors are located (Figure 4D). We observed specific gains and losses of chromosome arms regionalized on the UMAP in a cluster-specific manner (Figure 4E-H, Figure S4C). Consistent with published data, the aggressive region of the UMAP (cluster A) shows loss of 1p, 6q, 10q and 14q (Figure 4E-H)^24^. We also observed loss of 9p, 18p, 18q and gain of 17q enriched in the most aggressive regions (Figure 4I, S4I). Chromosome 1p loss and 1q gain was specific to cluster A, while gain of 1p was seen frequently in the rest of the UMAP (Figure 4E, 4I, S4G). Other groups have demonstrated PTEN mutations in aggressive tumors^3^. We show loss of 10q, which harbors PTEN, and low expression of PTEN in the most aggressive region (subcluster A3) (Figure 4H, S4H). Moreover, within cluster A, where the most aggressive tumors are, patients with loss of either 1p, 6q, 10q, or 14q were all associated with shorter time to recurrence than the patients without those CNAs (Figure 4J, 4K, Figure S4D-F).

### Meningioma subtypes frequently show expression patterns of developmental cell types

The meningioma UMAP reference landscape was generated using RNA-Seq data and therefore presents a great advantage of performing differential gene expression analysis and deciphering the underlying biology across meningioma subtypes. We first determined the differentially expressed genes in each cluster relative to the rest of the meningiomas. We then performed gene ontology (GO) analysis on Enrichr and the most prevalent GO terms in each cluster were used to discern the underlying biological signature for each of them (Figure 5A, 5B, 5C, Figure S5A, Table S5A)^40^. Cluster A was enriched for cell cycle, skeletal and cardiac muscle development, and DNA replication and repair while cluster B was specific to immune cells and function (Figure 5A, 5B) (see Figure 5 in Oncoscape). Although cluster C had relatively fewer GO terms that did not point towards a specific biological signature, *SMO* mutations were enriched in cluster C (Figure S5A, S5B). Accordingly, regulation of Smoothened signaling and SHH pathway were upregulated within cluster C (Figures S5A, S5C). Similarly, cluster D had a broader collection of GO terms, however, enriched for *AKT1* mutations (Figure S5A, S1F). Clusters E and F enriched for epidermis development and vascular development respectively (Figure 5C, Figure S5A). It is worth noting that KLF4, a transcription factor involved in skin development, was highly mutated in cluster E tumors—specifically, the K409Q mutation (Figure S1F)^41^. Cluster G, which had one of the worst outcomes, was enriched for neuronal functions including neurotransmitter / synaptic transmission and nervous system development. Cluster H and I were enriched for protein translation, macromolecule biosynthesis and mitochondrial functions, however, because they contain only Hong Kong patients, a larger sample size is necessary to confirm the identified biological signatures.

**Figure 5.**
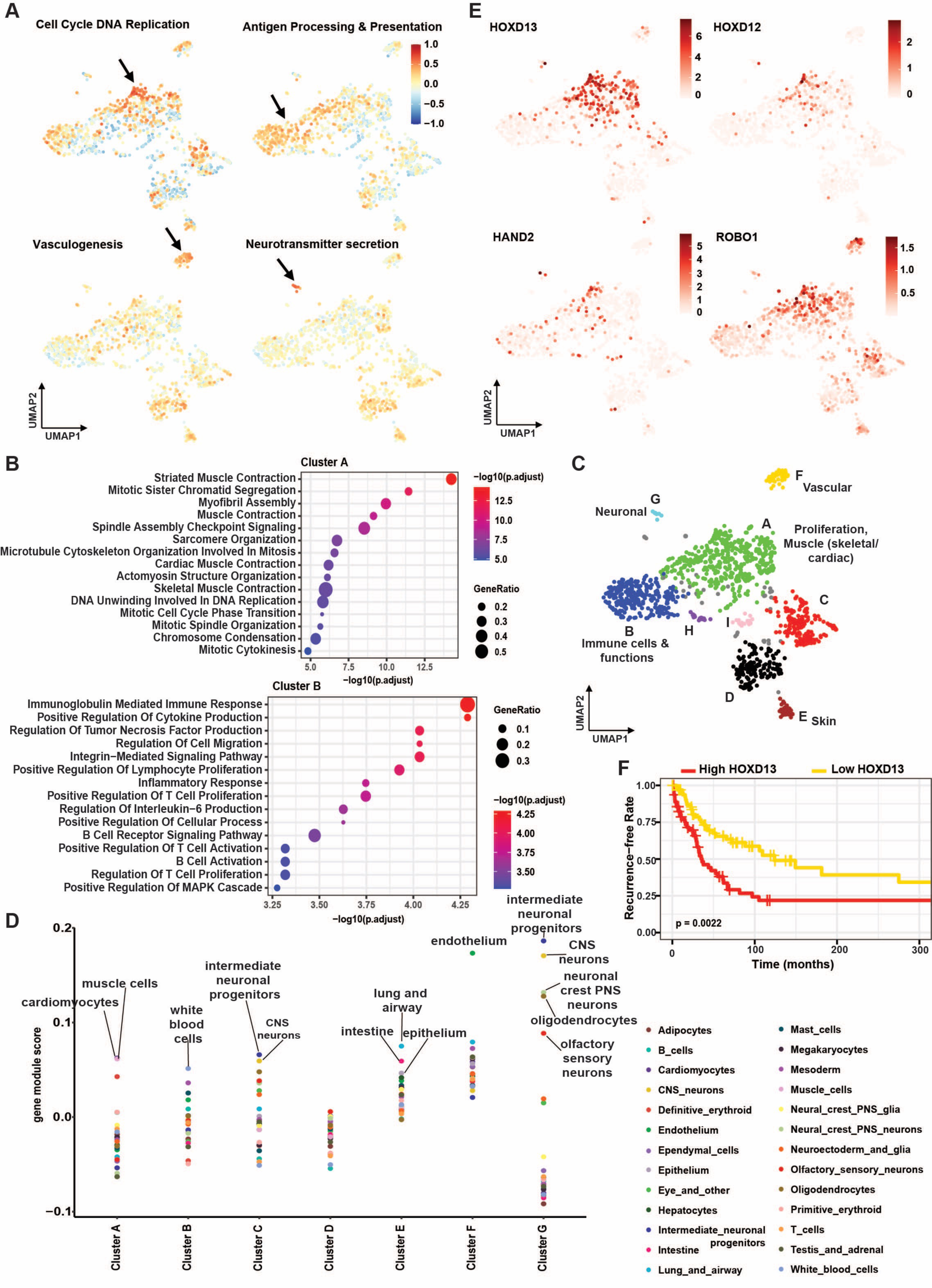
Biological significance of meningioma subtypes. (A) Visualization of GSVA scores across the UMAP for selected Gene Ontology Biological Processes (GO BP) terms. A score closer to 1 suggests upregulation of the respective gene set while a score closer to −1 suggests downregulation of the respective dgene set. (B) Top 15 GO BP terms enriched in clusters A and B (C) Summary of biological significance of each cluster (D) Mouse embryonic cell types enriched in each cluster (top hits). Muscle cells and cardiomyocytes in cluster A, white blood cells in cluster B, intermediate neuronal progenitors and CNS neuron in cluster C, lung and airway, intestine and epithelium cells in cluster E, endothelium cells in cluster F and several neuronal cells in cluster G were significantly enriched (Welch’s two sample t-test; p.value < 2.2e-6) (E) Visualization of gene expression profiles for genes known to be involved in embryonic limb development. (F) Kaplan Miere plots showing high recurrence-free rate in tumors with low HOXD13 levels and low recurrence-free rate in tumors with high HOXD13 levels (p.value = 0.0022). (see Figure 5 in Oncoscape) See also Figure S5, Table S5A and Table S5B.

Biological signatures that we identified were in line with tumor classifications based on methylation profiles presented by other groups. Tumors classified by UCSF and University of Toronto groups as highly proliferative were enriched within cluster A of this UMAP. Immunogenic tumors of their classification overlapped with cluster B (Figures 2B, 2C).

Some of these clusters were notably enriched with developmental GO terms, indicating potential parallels between the biology of these tumors and embryonic development. Therefore, to further learn about the potential association of meningioma subtypes with developmental cell types, we compared the identified cluster-specific gene signatures to mouse embryonic cell types. We leveraged the transcription profiles of a series of mouse embryonic developmental stages and hundreds of cell types put together by the Shendure lab^42^. In line with what GO terms suggested, we found that cluster A was enriched specifically for muscle progenitor cells and cardiomyocytes while cluster B was enriched for immune cells (Figure 5D, Figures S5D, Table S5B). Cluster C was enriched for neuronal cells; however, cluster D was not enriched for any specific embryonic cell type (Figure 5D, Figures S5D, Table S5B). Consistent with GO terms, epithelial cells and endothelial cells were top hits in clusters E (skin-related) and F (vascular-related) respectively (Figure 5D, Figure S5D, Table S5B). Similar to what GO terms suggested, Cluster G was enriched for various neuronal cells (Figure 5D, Figure S5D, Table S5B).

Cluster A, where the GO terms suggested muscle development, exhibited enrichment of *HOXD12/13*, *Hand2*, and *Robo1* genes that were highly expressed in the subcluster A3, which had one of the worst outcomes (Figure 5E). Relative enrichment of expression of these genes suggests the possibility that the underlying biology of this cluster may resemble embryonic limb development^43^. Alternatively, biology of this cluster could be related to the entirety of embryonic state in which the limb is developing rather than limb development specifically. Interestingly, high *HOXD13*-expressing tumors in cluster A had a significantly shorter time to recurrence than the low *HOXD13*-expressing tumors of that same cluster which may further suggest that limb-related development correlates with the most common aggressive meningiomas (Figure 5F).

### Recurrent tumors remain largely in the cluster from which they arise

In our collection of tumors there were several cases where samples were resected from multiple tumors from the same patient. We identified three scenarios: recurred tumors, multiple individual tumors from different brain regions, and progressed tumors due to incomplete surgical resection. We evaluated their location on the UMAP to further understand how their biology and outcome might differ with time. Recurrent tumors remained within the clusters that they were found originally, and vectors (between two tumors of the same patient) do not point towards a more aggressive region of the map (Figure 6A, Table S6a). Regardless of the time between recurrences, this result suggests that the recurred tumors’ biology and outcome do not vastly differ from the initial tumor. Within the collection of tumors were four cases where different, multiple tumors had occurred in different brain regions (Figure 6B, Table S6b). Most of them were patients with *NF2* loss. While some of the tumors presented similar biology and outcome, some were vastly different from each other. Two cases where the tumors progressed due to previous incomplete surgical resection mapped within the same cluster. (Figure 6C, Table S6c).

**Figure 6.**
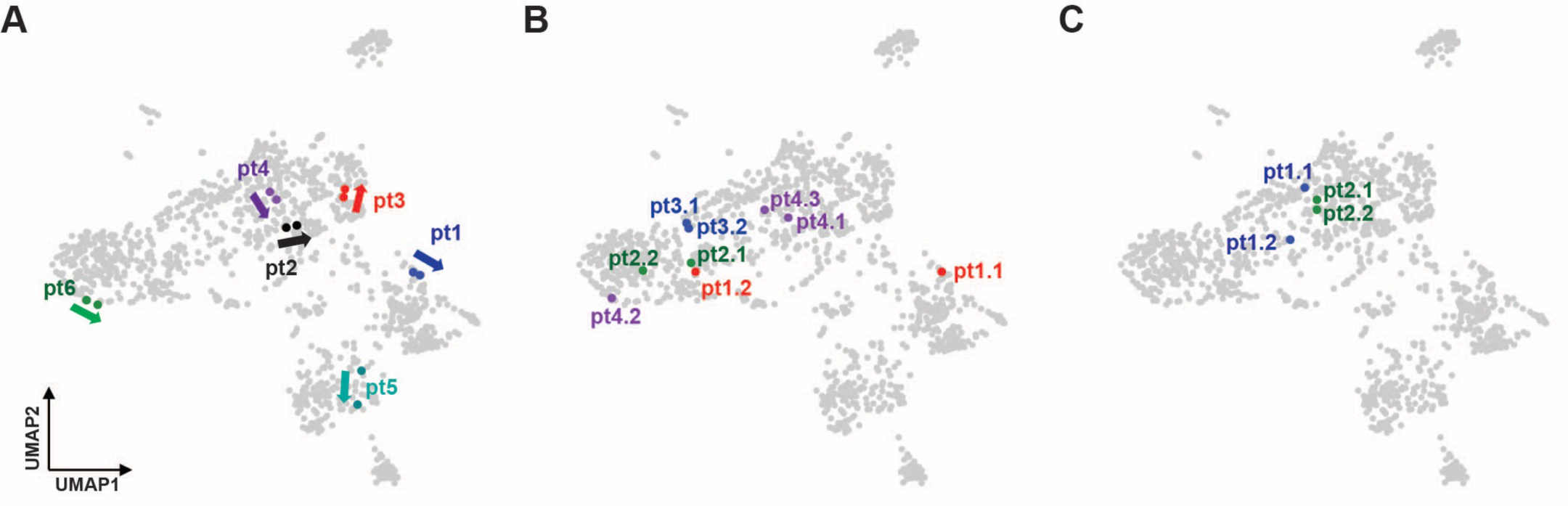
Evolution of multiple tumors from the same patient. (A) Primary and recurred tumors from the same patient are mapped. Arrows shows the direction from the 1^st^ tumor to the 2^nd^ tumor of a specific patient. Tumors from a single patient is distinguished by the colors (pt = patient). (B) Multiple individual tumors occurred within the same patient. Each patient distinguished by different colors (e.g., pt1.1 = patient 1 tumor 1, pt1.2 = patient 1 tumor 2). (C) Primary and progressed tumors (e.g., pt1.1 = patient 1 tumor 1, pt1.2 patient 1 tumor 2 (progressed). See also Table S6.

### Overlaying new patients onto an existing reference UMAP

The above data have shown that the biology and outcomes of meningiomas are regionally located in our UMAP reference landscape. Therefore, the nearest neighbors on the UMAP to a given tumor can serve as references from which a tumor’s biology and likely outcome can be inferred. However, for us to make such inferences, we must be able to reliably map a new patient onto our reference UMAP. To this end, we developed and validated an algorithm that accurately places new patients on our reference map.

Our placement method uses a weighted, nearest-neighbors approach that leverages an ensemble of UMAP models (Methods). Briefly, we pre-trained 100 UMAP models with different initializations on our reference dataset (Figure 7A) and used each model to map a new patient to a distinct two-dimensional embedding (Figure 7B). We then used the location of the new patient in each embedding to determine which reference samples are the 100 nearest neighbors of the new patient in each embedding within a radius determined using cross-validation (Figure 7C).

**Figure 7.**
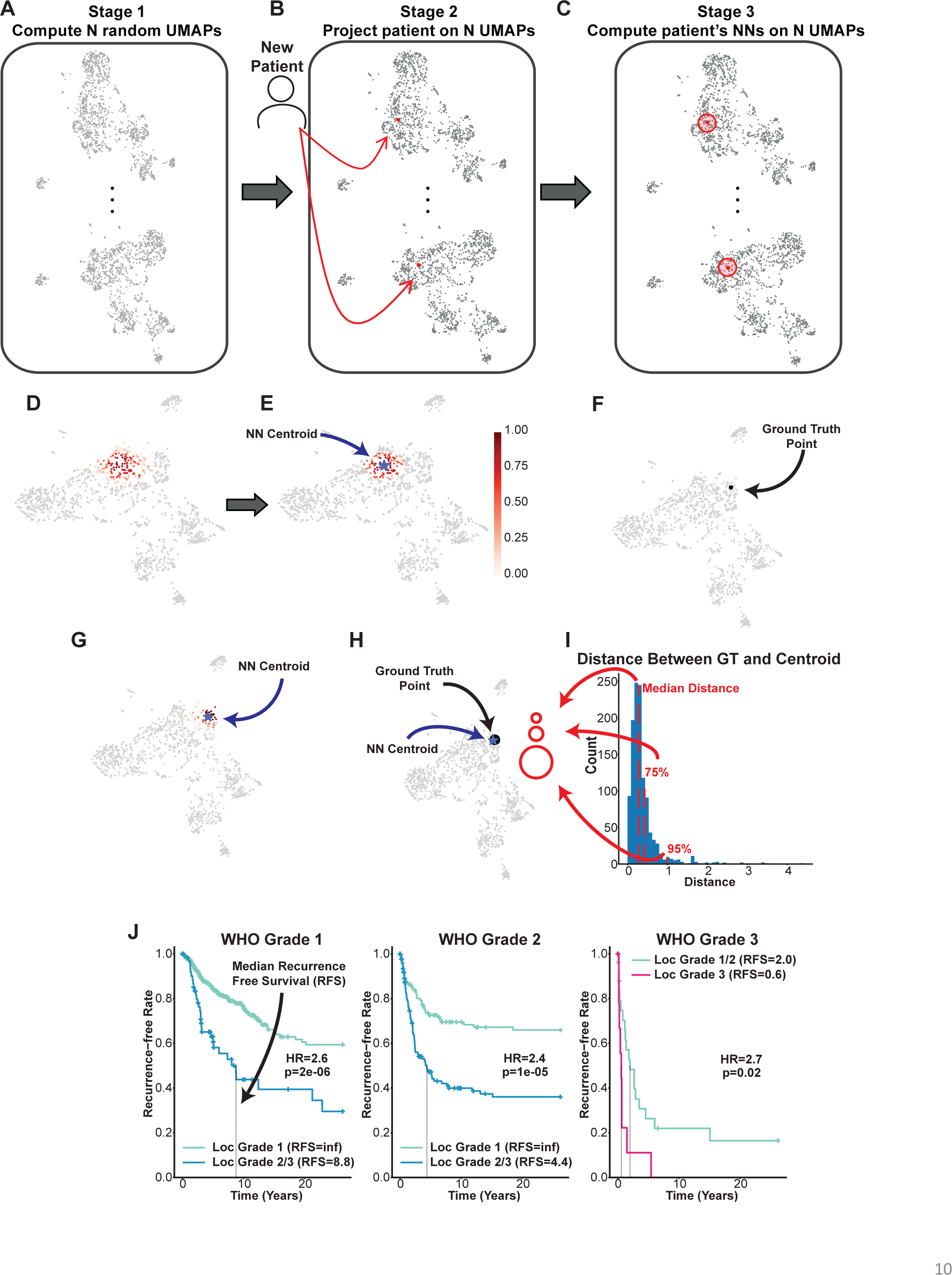
Overlaying new patients on to the reference UMAP. (A) Two of 100 UMAP embeddings produced by 100 pre-trained UMAP models trained with different random states. (B) New patient TPM data is mapped onto all 100 UMAP embeddings using the pre-trained UMAP models. (C) For each UMAP embedding, the nearest 100 neighbors are chosen subject to a radius R determined by cross-validation. (D) Example plot of the reference UMAP with samples colored by the frequency each reference sample in our reference dataset is a nearest neighbor of a new patient. (E) Illustration of the placement of a new patient at the centroid of the nearest neighbors weighted by the frequency vector in (D) after outlier exclusion. (F) The ground truth location of a reference sample during cross-validation. (G) The placement of a reference sample using our placement method during cross-validation. (H) Comparison of the ground truth placement of a reference sample and the centroid it is mapped to during cross-validation. (I) The distribution of the distances between the ground truth placement of a reference sample and its centroid placement for all reference samples during cross-validation. (J) Kaplan-Meier curves for location grade predictions within WHO grade 1, 2, and 3 meningiomas in our reference dataset. See also Figure S7.

This results in 100 sets of nearest neighbors from the reference dataset. This information is used to determine how frequently each reference sample in our reference dataset is a nearest neighbor of the new patient (Figure S7A). Finally, using this frequency information and the coordinates of the reference samples on our reference UMAP, we computed the centroid of the coordinates of the reference samples on our reference UMAP weighted by the frequency with which these samples were nearest neighbors of the new patient (Methods) (Figure 7D, 7E). We used this centroid as the final placement location of a new patient on our reference UMAP.

To establish the reliability of our placement algorithm, we used cross-validation to assess how far samples in the reference dataset moved when they were removed from the dataset and mapped back onto the reference UMAP. First, we considered each reference sample’s location in the reference UMAP as ground truth (Figure 7F). Next, we iteratively removed each sample, retrained our UMAP models without that sample, and used our placement method to place each sample back onto the reference map (Methods) (Figure 7G). Last, we computed the Euclidean distance between each reference sample’s ground truth position and its predicted location (Figure 7H). Nearly all reference points were mapped within a small radius of their true location (Figure 7I). The reliability of our placement method was also confirmed by evaluating its predictive power. Cross-validated results showed that our method was able to predict patient cluster membership accurately (AUC=0.98) by simply predicting the cluster most common in the samples around which the patient was placed (Figure S7B-S7D, Methods).

Cross-validated results also demonstrated the potential prognostic utility of our reference landscape. To leverage a patient’s location on our reference map, we assigned a location-based tumor grade to each sample in our reference dataset that corresponded to the WHO grade most common in the sample’s surrounding samples once remapped onto our reference map (Methods). Our results indicated that our predicted location grade was a superior risk indicator compared to WHO grade within WHO grade 1 and WHO grade 2 meningiomas and, to a lesser extent, WHO grade 3 tumors. In univariate analyses, WHO grade 1 tumors were separated into predicted location grade 1 and predicted location grade 2/3 tumors with dramatically different recurrence-free survival (HR=2.6, p=2e-06); similarly, WHO grade 2 tumors classified as location grade 1 had significantly better recurrence-free survival compared to tumors classified as location grade 2 or 3 (HR=2.4, p=1e-05) (Figure 7J, Figure S7E). Additionally, among WHO grade 1 and 2 tumors, a multivariate analysis shows our predicted location grade is an independent predictor of recurrence-free survival compared to WHO grade in our reference dataset (Figure S7F). However, although WHO grade 3 tumors classified as location grades 1 or 2 had more favorable outcomes than those predicted to be location grade 3 (HR=2.7, p=0.02), all WHO grade 3 tumors experience short times to recurrence. Thus, despite the prognostic power of our UMAP landscape, histopathology plays a crucial role in assessing patient risk. We do not propose this location-based tumor grade as an alternative to current classification systems; rather, we present these predictions to highlight the predictive power of the reference landscape. The ability to place new patients on this landscape makes our findings relevant for clinical applications, and we believe this study lays a strong foundation to build a reliable predictive tool in the future.

## Discussion

Most studies thus far have categorized meningiomas as malignant or benign. There are several classification systems currently available that are based on histomorphology of tumors and a limited number of genetic alterations. In this study, our goal was to determine the complexity of the meningioma population and decipher the underlying biology of these tumors using a comprehensive transcriptomic-based approach. We identified a complex landscape of multiple meningioma subtypes comprising of nine general map regions and determined that there is more than one aggressive type (clusters A, C and G). The reference UMAP regionalized meningioma subtypes based on *NF2* status (*NF2* wild-type and *NF2* mutant) and underlying biology (as proliferative and immunogenic) that are in line with previous studies^3,4,21^. More importantly, our study further extends the understanding of meningioma subtypes by adding a granular classification such as subtypes related to muscle development, skin, vascular and neuronal development, and separate clusters within the *NF2* wild-type region (clusters C and D, which enrich for *SMO* and *AKT1* mutations, respectively). In addition to GO term analysis, we compared clusters against different cell types during mouse embryonic development, where we show that the biology of clusters depicts both pathways and developmental cell types. In summary, we designated the subtypes as proliferative and muscle development (cluster A), immunogenic (cluster B), benign *NF2*-wild-type with *SMO* mutations (clusters C), benign *NF2*-wildtype with *AKT1* mutations (cluster D), skin (cluster E with *KLF4* mutations), vascular (cluster F) and neuronal (cluster G). Further studies are necessary to decipher the biology of clusters C, D, H, and I.

Within main clusters, we identified subregions that perform differently in terms of patient outcome (aggressive regions A3, A4, C2, and G). These subregions were further ascertained by the UMAP colored in by UCSF gene signature (ratio of upregulated/downregulated genes: Figure 2F). Moreover, we observed CNA patterns that highlighted subregions within clusters. For example, chr 10p and 10q in subcluster A3; chr 5q, 19p and 19q in subclusters B1/B2; and chr 1p, 19p and 19q in subclusters F1/F2. Identification of such subregions underscores the importance of predicting the outcome of a new patient based on nearest neighbors within a subregion instead of overseeing it as a whole cluster. The addition of more samples to the map will likely allow delineation of these subregions with higher accuracy.

Our dataset consists of 33 pediatric (< 18 yrs) and 49 young adults (19 - 30 yrs). The majority of these young patients mapped onto the region between cluster A and B. Consistent with other tumor types, pediatric patients and young adults harbored more gene fusions as opposed to copy number alterations (data not shown). YAP1-MAML2 fusion was more prominent among younger patients while NF2 fusions were enriched among adults. Further investigation is warranted comparing fusions in adults and young patients with a larger cohort of the latter. Furthermore, we observed a large fraction of fusions with non-coding genes. Although we limited our current study to fusions involved with protein coding genes, it would be worth deciphering the role of non-coding fusions in meningiomas in future studies.

In addition to identifying meningioma subtypes and understanding their biology, we developed a method to overlay prospective patients on our reference UMAP to infer their biology and clinical outcomes from their nearest neighbors. We used cross-validation to verify that patients were accurately placed on the reference UMAP both by measuring the distance our method places reference samples from their location in the reference UMAP and by assessing how well our method predicts cluster membership. In addition to cluster membership, we also used our method to assign patients a location grade based on the distribution of the WHO grade of their surrounding samples. Some tumors classified as benign and WHO grade 1 were located in the most aggressive region of our UMAP reference landscape, which suggests that despite the histopathological grading their underlying biology and outcome are similar to more malignant tumors. Therefore, our transcriptomic-based UMAP landscape of meningioma may provide a better understanding of the patient biology and outcome. If clinical data was associated with the patients of the reference landscape, the nearest neighbor analysis could also be used to identify what therapeutic approaches were successful in the patients with tumors most similar. In the case of meningiomas specifically, there are no targeted therapies that work in a subset of these tumors. However, in other tumor types, this kind of analysis could help distinguish those likely to respond based on the underlying biology of the disease type.

We combined multiple datasets from various sources, corrected for batch effects, and generated the largest meningioma reference landscape to date with comprehensive analysis of all protein coding genes. One of the main goals of our reference landscape was to better understand the underlying biology of meningioma subtypes and using transcriptomic data benefited us with the ability to extract biological information such as expression of specific genes and pathways in a straightforward manner. We defined multiple meningioma subtypes that predicts tumor biology and patient outcome. To our knowledge, this is the first paper to put forth a reference map of a disease on an interactive online tool. Oncoscape is not only an attractive visualization platform for the figure panels, it also provides the opportunity for multifaceted exploitation of the map while mining for various metadata such as patients’ clinical information and genome-wide gene expression, gene fusions, copy number alterations and biological pathways. Overall, this study shows how we can harness the power of combining multiple datasets to extract further biological information of a particular disease. This approach may be useful in other tumor types, there is no reason to believe that success of this analysis will be unique to meningiomas.

## Acknowledgments

We thank members of the Holland lab at Fred Hutch Cancer Center for valuable discussions and collaborators for sharing their data and metadata. We thank the Fred Hutch core services: Specimen Processing & Research Cell Bank and, Genomics and Bioinformatics Services. This research was supported by funding from the Fred Hutch Cancer Center (E.C.H), 1R35 CA253119-01A1 (E.C.H), P30 CA015704 (Genomics & Bioinformatics Shared Resource <u>RRID:SCR_022606</u> of the Fred Hutch/University of Washington/Seattle Children’s Cancer Consortium).

## Author contributions

Conceptualization, H.N.T, D.A.B, N.N., S.A., F.S., and E.C.H.; Methodology, H.N.T, D.A.B., N.N., S.A., F.S., and E.C.H.; Formal Analysis, H.N.T., S.A. and N.N.; Software, M.J.; Investigation, H.N.T., D.A.B., N.N., S.A., C.P., M.F. and C.Q.; Resources, C.W.E, W.C.C, P.S., F.N., J.W., T.J.K., K.D.A., A.J.P., G.Z., C.P., F.S., M.F. and D.R.R.; Data Curation, H.N.T, D.A.B., S.A. and N.N., Writing – Original Draft, H.N.T., N.N., and E.C.H; Writing – Review & Editing, S.A., F.S., P.J.C., D.R.R., J.S. and E.C.H; Visualization, H.N.T., N.N. and M.J.; Supervision, J.S., M.F. and E.C.H., Funding Acquisition, E.C.H.

## Methods

### Key resources

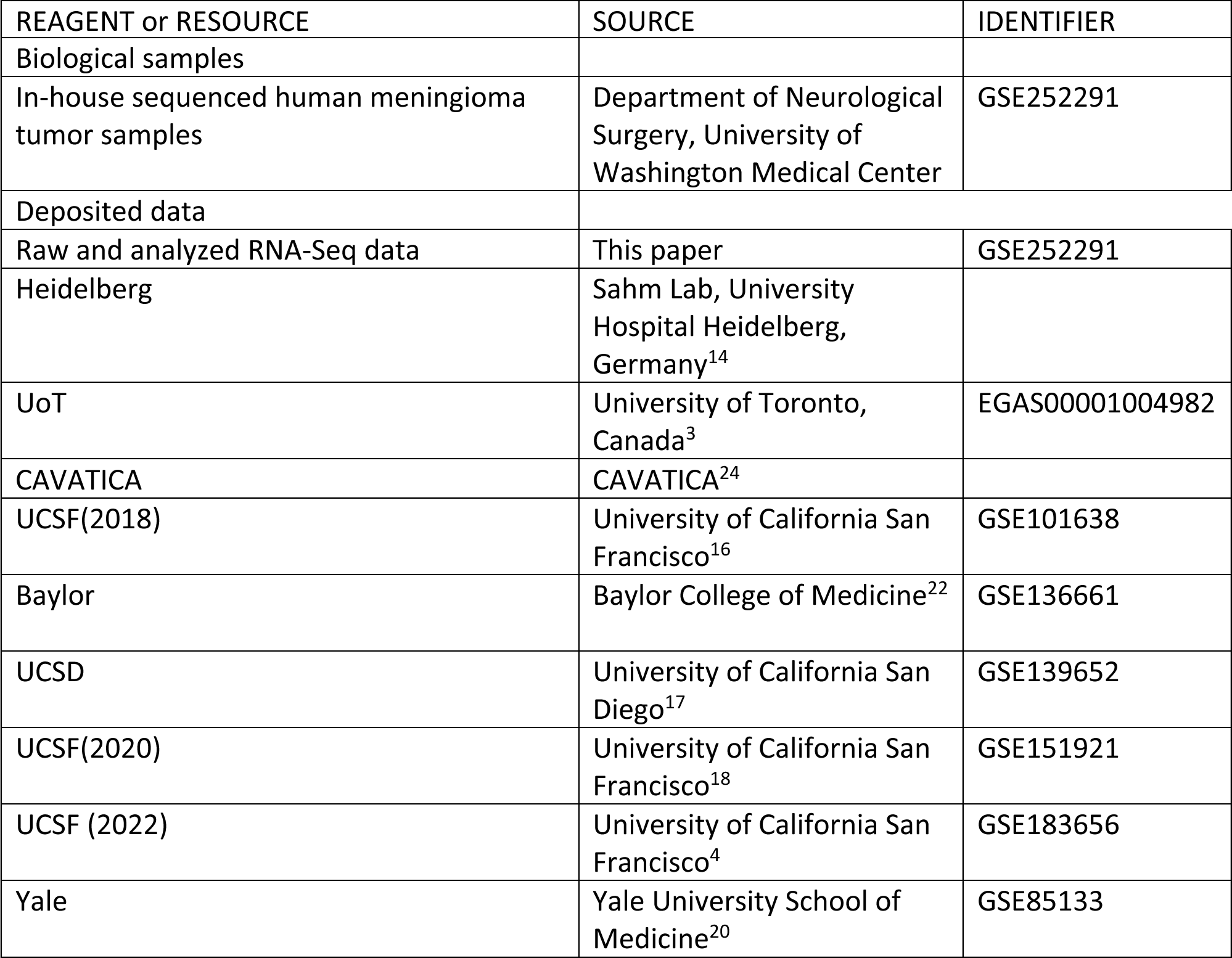
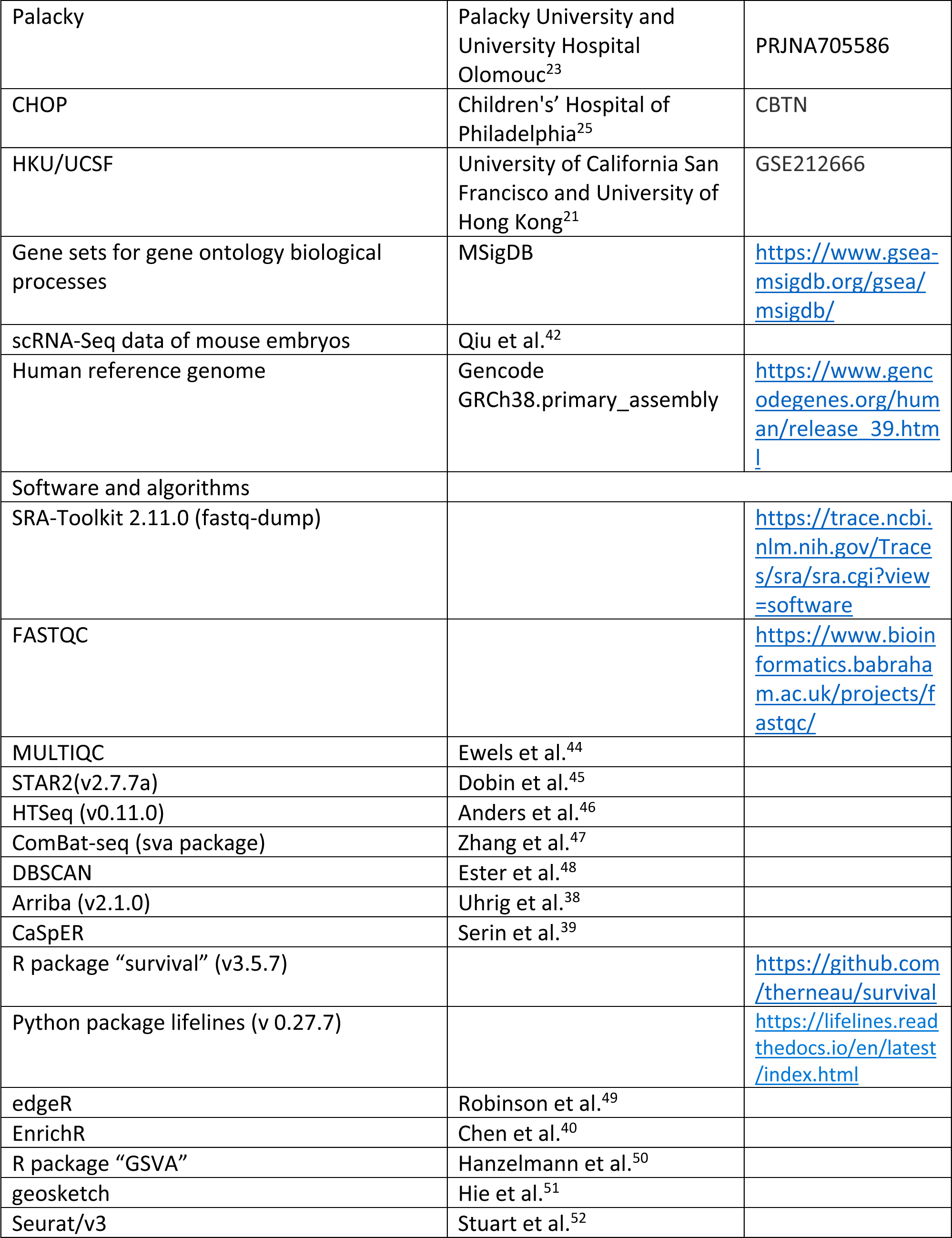

### Collection of specimens and clinical data from University of Washington

Studies were conducted in accordance with the U.S. Common Rule ethical guidelines. Tissue was collected from study participants admitted at the University of Washington Medical Center, Department of Neurological Surgery. The respective clinical data was extracted from the University of Washington, School of Medicine clinical database. Data and specimen collection were reviewed and approved by the University of Washington Institutional Review Board and Human Subjects Division. Written-informed consent was obtained from all subjects. Patients underwent surgery at the University of Washington Medical Center between January 1, 1998 and June 1, 2023. Samples were collected and stored in −80°C. Clinical data were gathered regarding history, demographics, imaging, neuropathology reports, operative information, adjuvant treatment, and outcomes. Resected tumors were graded according to the current criteria (DOI: 10.1093/neuonc/noab106) and correlated with clinical information, when advised. Histologic subtype, mitoses, Ki-67/MIB, sheeting, macronuclei, hypercellularity, invasion, necrosis, TERT promoter mutations (DOI: 10.1093/jnci/djv377)) and CDKN2a/b homozygous deletions (DOI: <u>10.1007/s00401-020-02188-w</u>) were recorded. Specimens were reviewed by three neuropathologists and neurosurgeons. Total resection was defined as absence of residual enhancement on postoperative MRI within 48 hours of surgery. Recurrence was defined as at least 1 cm of enhancement on subsequent MRI. Progression was considered to be at least 1 cm of growth of residual tumors detected on MRI after surgery.

### Specimen processing for RNA-Seq

RNA was extracted using the Qiagen RNeasy Plus mini kit. Total RNA integrity was checked using an Agilent 4200 TapeStation (Agilent Technologies, Inc., Santa Clara, CA) and quantified using a Trinean DropSense96 spectrophotometer (Caliper Life Sciences, Hopkinton, MA). Samples with RIN < 5 were removed from further analysis. RNA was normalized to 50ng/ul and 500ng was submitted for library preparation. RNA-seq libraries were prepared from total RNA using the TruSeq Stranded mRNA kit (Illumina, Inc., San Diego, CA, USA). Sequencing was performed using an Illumina NovaSeq 6000 employing a paired-end, 50 base read length (PE50) sequencing strategy. RNA extraction was performed by the Specimen Processing & Research Cell Bank at Fred Hutch. Library preparation and sequencing was performed by the Genomics and Bioinformatics Core Services at Fred Hutch.

### Collection of publicly available RNA Sequencing data

Raw sequencing data of human meningioma samples were obtained from respective public data repositories (Table S1) except Heidelberg dataset which was obtained from the data repository of the Dept. of Neuropathology at the University Hospital Heidelberg. SRA files downloaded from GEO were converted to fastq files using fastq-dump from the SRA-Toolkit (v2.11.0).

### RNA-Seq data processing and visualization

Quality check on raw RNA sequencing data was done using FastQC (v0.11.9) (https://www.bioinformatics.babraham.ac.uk/projects/fastqc/) and MultiQC (v1.9) tools^44^. RNA sequencing reads were aligned to the Gencode GRCh38.primary_assembly genome using STAR2 (v2.7.7a) and then using HTSeq (v0.11.0) reads were counted for each associated gene using the Gencode V39 primary assembly annotations^45,46^. Raw gene counts from each dataset were combined and corrected for batch effects using ComBat-seq from the R package “sva”^47^. Gene expression values from combined datasets were normalized and converted to units of log2 transcripts per million (log2(TPM+1))^26^. Uniform Manifold Approximation and Projection (UMAP), a dimensionality reduction method, was applied on normalized counts from 19979 protein-coding genes to create the meningioma reference landscape^27^. UMAPs were constructed using the R package “umap” (https://cran.r-project.org/web/packages/umap/index.html).

### Clustering using DBSCAN

DBSCAN (density-based spatial clustering of applications with noise) was used to confirm the clusters identified by UMAP^48^.

### Obtaining gene fusion using RNA-Seq

Arriba (v2.1.0) was used to compute gene fusions from two-pass STAR-aligned RNA-Seq reads^38^. All fusion analyses were restricted to fusion calls Arriba indicated were high confidence. Using gencode.v38.annotation.gtf.gz from hg38 release 44 (GRCh38.p14) from https://www.gencodegenes.org/human/ we determined whether a gene was protein coding and selected only fusions with at least one coding gene involved. Fusions that recur at least twice within the dataset were used to calculate fusion burden.

### Obtaining Copy Number Alternations (CNA) using RNA-Seq

Large scale / chromosome arm level copy number alternations were estimated for all tumors using the P package CaSpER (https://rpubs.com/akdes/673120) on bulk RNA-Seq data^39^. BAFExtract source code, genome list and genome pileup directory were downloaded from https://github.com/akdess/. hg38 cytoband and centromere information were downloaded from UCSC. GTEX RNA-Seq data from normal frontal cortex and hippocampus (dbGaP Accession: phs000424.v8.p2) were used as control samples in the CaSpER analysis (GTEX sample IDs: SRR1147618, SRR1334440, SRR1337431, SRR1342045, SRR1348360, SRR1354446, SRR1360128, SRR1375571, SRR1388305, SRR1402900, SRR1408368, SRR1413562, SRR1416477, SRR1435775, SRR1453341, SRR1471817, SRR1488367, SRR1488651, SRR1500868).

### Kaplan-Meier curves

Kaplan-Meier curves were generated using the information on time to recurrence of each sample. To perform the calculations, we only selected tumors with known recurrence status (recurrence = yes/no) and known time to recurrence or last follow up. For tumors that were confirmed as non-recurrent however with no last follow up date, a default of 315 months was used as “Months of No Recurrence”. Kaplan Meier curves were plotted and p.values were calculated using the R package “survival” (v3.5.7).

### Differential gene expression analysis

To determine the biological signature of each cluster, differential expression analysis was performed between each cluster and the rest of the UMAP samples using edgeR^49^. Upregulated genes in each cluster were identified based on FDR (< 0.05) and log fold change (> 0.6) or fold change of 1.5 cut-off.

### Pathway analysis

Top 500 significant DE genes of each cluster (with the exception of cluster C = ∼700 genes) were analyzed using Enrichr^40^. Gene Ontology Biological Processes (GO BP) terms that were statistically significant (adjusted p value < 0.05) were manually curated to remove GO terms that had gene sets >80% overlapping. Dot plots were generated using ggplot2 (top 15 GO terms included). Complete lists of GO terms were attached as supplement information.

### GSVA pathway analysis

Gene sets for pathways from Gene Ontology Biological Processes were downloaded from Molecular Signature Databases (MSigDB) (v7.2) (https://www.gsea-msigdb.org/gsea/msigdb/collections.jsp). Gene Set Variation Analysis (GSVA) was performed on batch corrected log2(TPM) counts from all 1298 samples. GSVA scores obtained from 1 and −1 for each sample were visualized using ggplot2. Similarly, GSVA analysis was performed using UCSF gene set (34 genes upregulated or downregulated in aggressive meningioma). Upregulated and downregulated genes were considered as two separate gene sets.

### Embryonic cell type analysis

The scRNA-seq data of mouse embryos was downloaded from Qiu *et al.*^42^. In the study, over 11 million cells were profiled from mouse embryos during organogenesis and fetal development, with every 6 hours temporal resolution ranging from embryonic day 8 (E8) to postnatal day 0 (P0). This resulted in the identification of ∼190 cell types. To save computational time and memory, the dataset was downsampled to ∼1 million cells using geosketch^51^. Top 500 upregulated protein-coding genes in each cluster (with the exception of cluster C = ∼700 genes) were obtained from the previously mentioned differential expression analysis. A gene module score was calculated for individual groups of genes using the *AddModuleScore* function implemented in Seurat/v3^52^. Gene module score was calculated for each cell type and the mean score of each major cell cluster was calculated for Figure 5D. Welch two sample t-test was performed to statistically confirm the top cell clusters enriched in each meningioma cluster.

### Placing new patients on UMAP reference map

The stages of our algorithm, which places new patients on our UMAP reference map, are described below. These stages consist of pre-training UMAP models, mapping new patients to embeddings generated by pre-trained UMAP models, determining nearest neighbors on UMAP embeddings, aggregating all sets of UMAP embedding-derived nearest neighbors, and using the frequency of these nearest neighbors to compute a centroid. This last process also involves removing outliners before computing the centroid.

#### • UMAP pretraining

We train *K* = 100 UMAP models on the 1298 samples in our RNA-Seq reference dataset *D* using a different random state for each UMAP model. We denote each of the *K* pre-trained UMAP models by the function.

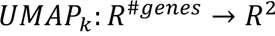

which maps any TPM sample to two dimensions. In this way, we can represent the embedding of the reference dataset given by each pre-trained UMAP model as *UMAP*_*k*_(*D*). We denote the *x*-coordinate and *y*-coordinate of every reference sample *p*_*i*_ in an embedding given by *UMAP*_*k*_ as *UMAP*_*k*_(*p*_*i*_)_1_ and *UMAP*_*k*_(*p*_*i*_)_2_, respectively. Similarly, we will refer to the *x*-coordinate and *y*-coordinate of every reference sample *p*_*i*_ in the reference landscape as *UMAP*(*p*_*i*_)_1_ and *UMAP*(*p*_*i*_)_2_. All UMAP embeddings given by *UMAP*_*k*_are normalized by centering the embeddings and scaling points so that the average distance between points is 1.

#### • Mapping samples to embeddings given by pre-trained UMAP models

Given *K* pre-trained UMAP models, we can place any new patient *P* onto a UMAP embedding *UMAP*_*k*_(*D*) by passing the TPM data of that patient (*P*_*TPM*_) through the UMAP model *UMAP*_*k*_. We denote this position in the embedding as *UMAP*_*k*_(*P*_*TPM*_), where we will refer to the *x* - coordinate of *UMAP*_*k*_(*P*_*TPM*_) as *UMAP*_*k*_(*P*_*TPM*_)_1_and *y*-coordinate as *UMAP*_*k*_(*P*_*TPM*_)_2_.

#### • Computing 100 sets of nearest neighbors from pre-trained UMAP embeddings

To get a set of nearest neighbors of a patient *P* on an embedding generated by a pre-trained UMAP model *UMAP*_*k*_, we first compute the distances between the position of the new patient *P* in the embedding and the position of all other reference samples *p*_*i*_. Thus, for every patient *p*_*i*_in the reference dataset and every pre-trained UMAP model *UMAP*_*k*_we compute the square of the Euclidian distances *d*_*k*_(*p*_*i*_, *P*) between *UMAP*_*k*_(*P*_*TPM*_) and the 1298 positions *UMAP*_*k*_(*p*_*i*_) on each of the 100 embeddings generated by the 100 *UMAP*_*k*_ maps. The square root of *d*_*k*_(*p*_*i*_, *P*) is not taken for efficiency. We define *d*_*k*_(*p*_*i*_, *P*) as follows:

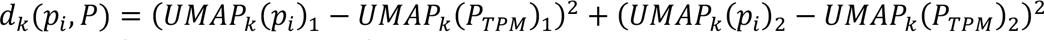

To obtain a list of nearest neighbors for the patient *P* on each UMAP model *UMAP*_*k*_, we choose the 100 samples *p*_*i*_ with the smallest *d*_*k*_(*p*_*i*_, *P*) under the constraint *d*_*k*_(*p*_*i*_, *P*) < *α*, where *α* = 0.05 is the square root of the Euclidian radius chosen so that the list of nearest neighbors would not span multiple unconnected clusters. We denote the set of nearest neighbors to a patient *P* on for a pre-trained UMAP model *UMAP*_*k*_as 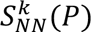.

#### • Computing the frequency vector used to weight the centroid calculation

We construct a matrix *M* whose values represent whether a sample in the reference dataset *p*_*i*_is a nearest neighbor of the new patient *P* in the *k*th UMAP embedding *UMAP*_*k*_(*D*). Formally, this can be expressed as

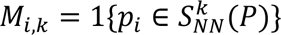

where 1 is the indicator function. To find the frequency vector *f*^*P*^of new patient *P*, we average the values of *M* across the *K* UMAP embeddings. This results in a vector of length 1298 whose value 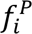 represents the frequency sample *p*_*i*_was a nearest neighbor of *P* across the *K* pre-trained UMAP embeddings.

#### • Compute centroid

The weighted centroid of a frequency vector *f*^*P*^ and the *x*-coordinate *UMAP*(*p*_*i*_)_1_ and *y* - coordinate *UMAP*(*p*_*i*_)_2_ of each sample on the reference UMAP is compute as follows:

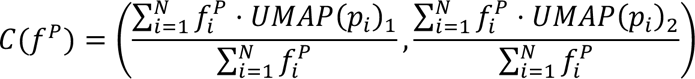

#### • Remove outliers in the frequency vector

To prevent samples *p*_*i*_ which may have been included in some set of nearest neighbors 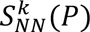 by chance, we create an adjusted frequency vector 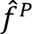 from *f^p^* by setting elements 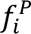 to zero for all such *p*_*i*_. We first set 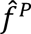 equal to *f*^*P*^. Next, we adjust 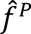 by setting values less than 0.25 to 0 and compute the weighted centroid 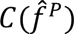 of 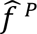 (described above). Then we compute the distance of all samples *p*_*i*_with non-zero 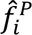 values to 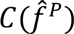 and remove those with distances greater than the 95% quantile. Last, we recompute the centroid 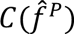 with the updated 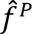 and set the values of 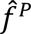 for samples farther than *R* = 0.75 from the centroid to zero. he radius *R* was chosen using cross-validated results and visual inspection.

#### • Final placement of the patient

To place a new patient *P* on our reference UMAP, we simply compute 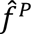 and place the patient at the coordinates given by 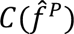 as described above.

#### • Uncertain placements

We compute a score to quantify the quality of our placements based on the distribution of frequencies in *f*^*P*^, which is computed as follows

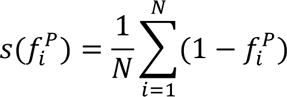

Here 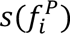 describes the average frequency every reference sample *p*_*i*_ is a nearest neighbor of *P* on the UMAP embeddings *UMAP*_*k*_(*D*) for samples that are nearest neighbors at least once (i.e., *p*_*i*_ such that 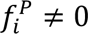). Lower values of 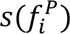 indicate that the sets of nearest neighbors were more consistent over all 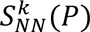 and suggest more reliable predictions. For this reason, we consider predictions for which 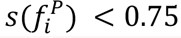. This threshold was established empirically using cross-validation.

#### • Cross-validation

To conduct cross-validation, we repeated the following procedure for each patient *p*_*i*_in our reference dataset *D*. First, we pre-trained 100 UMAP models on *D* ∖ *p*_*i*_, the reference dataset without the sample *p*_*i*_. Afterward, we treated *p*_*i*_ as a new patient and placed *p*_*i*_ on our reference UMAP at the location *C*(*f*^*pi*^) as described above. Finally, we computed the distributions of the Euclidean distances between the predicted centroid 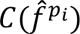 and the ground truth location of *p*_*i*_on the reference UMAP for each *p*_*i*_

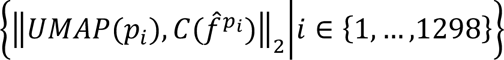

where

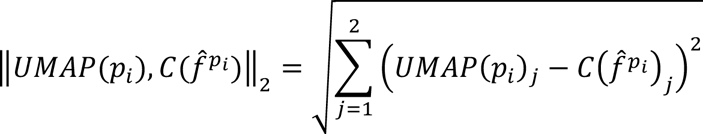

#### • Cluster prediction

To predict the cluster membership of a new patient, we mapped the patient onto our reference UMAP and computed its nearest neighbors.

We used a majority vote strategy which predicted the most common cluster found in this set of nearest neighbors. Cluster predictions given in the main text were computed via cross-validation.

#### • WHO grade prediction

We predicted WHO grade using the same procedure we used to predict cluster membership. We mapped a new patient onto our reference UMAP and computed its nearest neighbors. A majority vote strategy issued the most common WHO grade among the set of nearest neighbors as the WHO grade prediction. WHO grade predictions given in the main text were computed via cross-validation.

### Oncoscape integration

Matrix and clinical data were prepared for Oncoscape by converting them to cBioPortal formats (cbioportal.org). Custom settings, including colorings and precalculated views to match the paper’s figures, were stored in JSON in an Oncoscape updates.txt file. See https://github.com/FredHutch/OncoscapeV3/blob/master/docs/upload.md for details.

## Data analysis

All analysis including statistics and visualization were done in R version 4.2.0 (2022-04-22) as implemented in Rstudio (2022.05.999). Plots were generated using R basic graphics and ggplot2.

## Data availability

RNA-Seq raw data from meningiomas collected from UW/FHCC are deposited in GSE252291.

## Code availability

All custom code used in this study are available at https://github.com/FredHutch/MeningiomaLandscape-HollandLab

## Supplemental Figures and Tables Legends

**Figure S1.** Related to Figure 1. (A) UMAP generated without batch correction among datasets, colored in by datasets used in this study. (B) Principle Component Analysis (PCA) and t-distributed stochastic neighbor embedding (tSNE) analysis on non-batch corrected datasets. Tumors colored in by datasets. (C) PCA and tSNE on batch corrected datasets. Samples colored in by datasets (upper panels) and WHO grade (bottom panels). (D) Distribution of FFPE and Fresh Frozen tissue. (E) UMAP showing functional loss of *NF2*. Tumors with either *NF2* mutations or *NF2* fusions or Chr 22 loss are colored in blue (yes), tumors without any of those aberrations colored in yellow (no) and tumors without any data available colored in grey (na). (F) Tumors with *KLF4* (in red) and *AKT1* (in blue) mutations (G) Tumors with either one or more of *TRAF7, KLF4, AKT1 and SMO* mutations (H) Tumors colored in by time (months) to recurrence (I) Bar graph showing the percentage of males and females in Region 1 and in rest of the UMAP (All – Region1). ∼60% patients are male in Region 1 while ∼30% in rest of the UMAP. (J) Bar graph showing the percentage of patients’ age categories in Region 2 and in rest of the UMAP (All – Region2). Region 2 has a higher percentage of younger patients (1-30yrs). (K) Pie chart showing the total number of patients in each age category.

**Figure S2**. Related to Figure 2. (A) UMAP colored in by the Ferreira Grade available for a subset of tumors (B) UMAP generated using UCSF gene signature of 34 genes and colored in by WHO grade, recurrent status of tumors, Chr 22 loss and UCSF DNA Methylation group. (C) UMAP colored in by the Simpson Grade available for a subset of tumors.

**Figure S3**. Related to Figure 3. (A) UMAP colored in by clusters identified using k-means clustering method (B) UMAP colored in by clusters identified using hierarchical clustering (C) Subclusters of cluster B (B1 and B2) and their Kaplan Meier plots (p.value = 0.0003) (D) Subclusters of cluster F (F1 and F2) and their Kaplan Meier plots (p.value = 0.018).

**Figure S4**. Related to Figure 4. (A) NF2 gene fusions called by Arriba using RNA-Seq data. (B) Heatmap showing NF2 gene expression in tumors with chromosome 22q lost (−1), intact (0) or gained (1). (C) Loss (−1), gain (1) or intact (0) status of chromosome 22q in each tumor. (D, E, F) Kaplan-Meier plots showing recurrence-free rates of tumors with intact and deleted chr 10q (p.value < 0.0001), chr 22q (p.value = 0.16) and chr 14q (p.value < 0.0001). (G) Manhattan plots showing gains and losses of each chromosome arm across clusters D-I. (H) UMAP colored in by the expression of gene PTEN. (I) UMAPs colored in by the CNA status of each chromosome arm (gains (1) in red, losses (−1) in blue, and intact (0) in grey). Chr 13p, 14p, 15p, 21p and 22p are the short arms of acrocentric chromosomes, hence no CNA calls.

**Figure S5**. (A) Top 15 Gene Ontology Biological Processes (GO BP) terms enriched in clusters C-I. (B) Tumors that have only *SMO* mutations. (C) UMAP showing the enrichment of SHH pathway. (D) Gene module score calculated for genes upregulated in each meningioma cluster across ∼190 embryonic cell types (cell types grouped into 26 major clusters). Blue dash line marks the significantly enriched cell types in each cluster. See also Table S5B.

**Figure S7**. Related to Figure 7. (A) We average a matrix whose rows are samples in our reference dataset, whose columns are pre-trained UMAPs, and whose values indicate whether a particular sample in our reference dataset is a nearest neighbor of a new patient being placed on the reference UMAP. (B) Cross-validated results showed we were able to predict patient cluster method membership in sizable clusters (N>20) with an AUC over 0.98 by simply predicting the cluster most common in a patient’s nearest neighbors which surround the patient once placed on the reference map. (C) Confusion matrix showing the cross-validated results of cluster predictions. Clusters G, H, and I have fewer than 20 samples. (D) Many of the incorrect cluster predictions occurred on the cluster boundaries, where cluster membership was ambiguous, indicating our performance metrics may underestimate our classifier’s efficacy. (E) Kaplan Meier curves corresponding to location grade predictions. Location grades 1, 2 and 3 shown separately for each WHO grade. (F) A multivariate Cox proportional hazards model shows that location grade is a superior, independent prognostic indicator compared to WHO grade in WHO grade 1 and 2 tumors.

**Table S1. Related to Figure 1**. Thirteen bulk RNA-Seq datasets were combined to generate the meningioma UMAP.

**Table S4A. Related to Figure 4**. Gene fusions called by Arriba using bulk RNA-Seq data (high confidence fusions with at least one coding gene partner that recur at least twice within dataset).

**Table S4B. Related to Figure 4**. Copy number alterations called by CaSpER using bulk RNA-Seq data.

**Table S5A. Related to Figure 5**. Gene Ontology Biological Processes terms enriched within each cluster.

**Table S5B. Related to Figure 5**. Mouse embryonic cell clusters enriched in each meningioma cluster and a complete list of mouse embryonic cell types.

**Table S6. Related to Figure 6**. Multiple tumor samples from the same patient: recurrences, progressions, and multiple individual tumors.

