## Supplemental Figures and Tables for "Meningioma transcriptomic landscape demonstrates novel subtypes with regional associated biology and patient outcome"

**Figure S1. Related to Figure 1.**

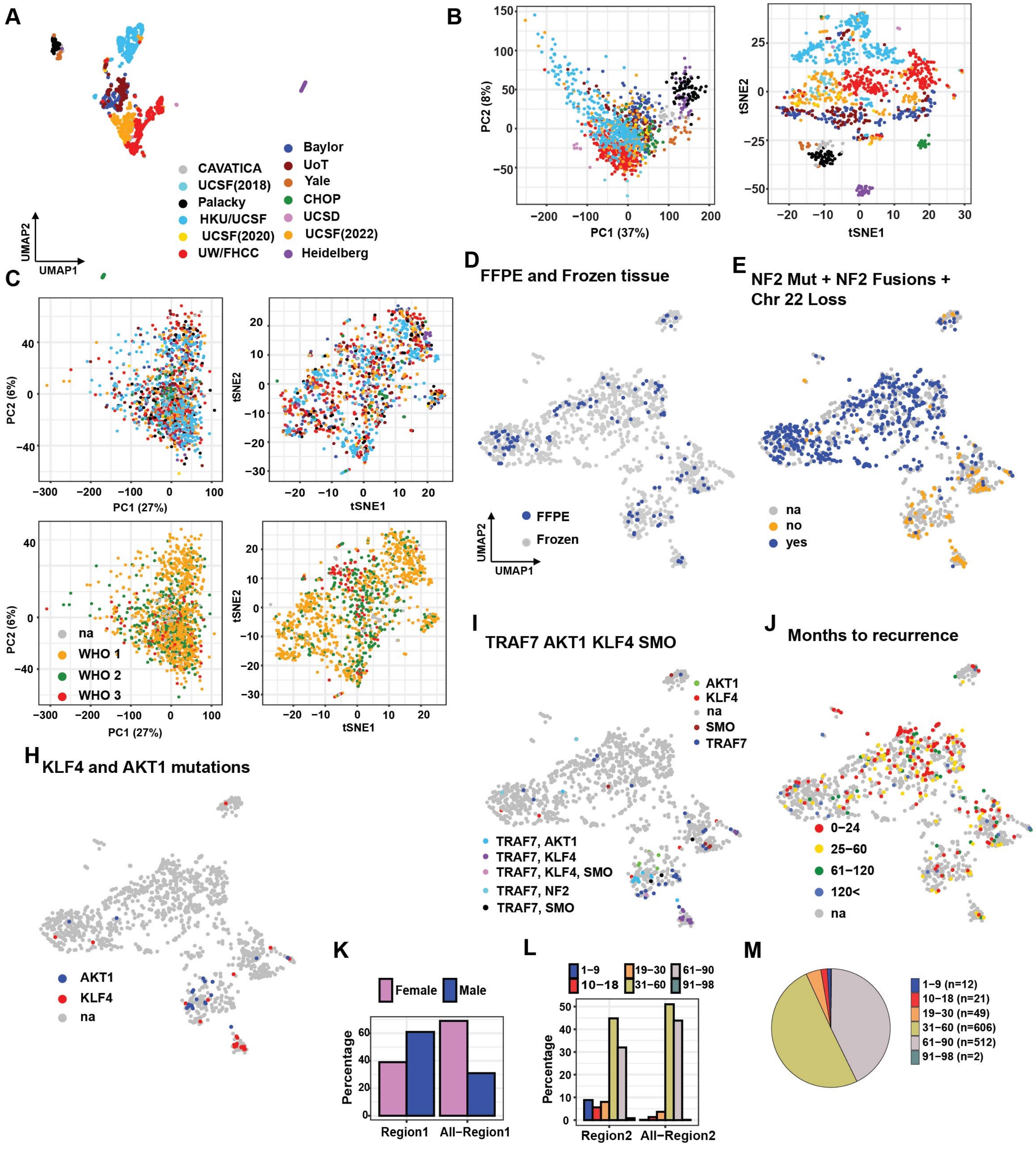

**Figure S2. Related to Figure 2.**

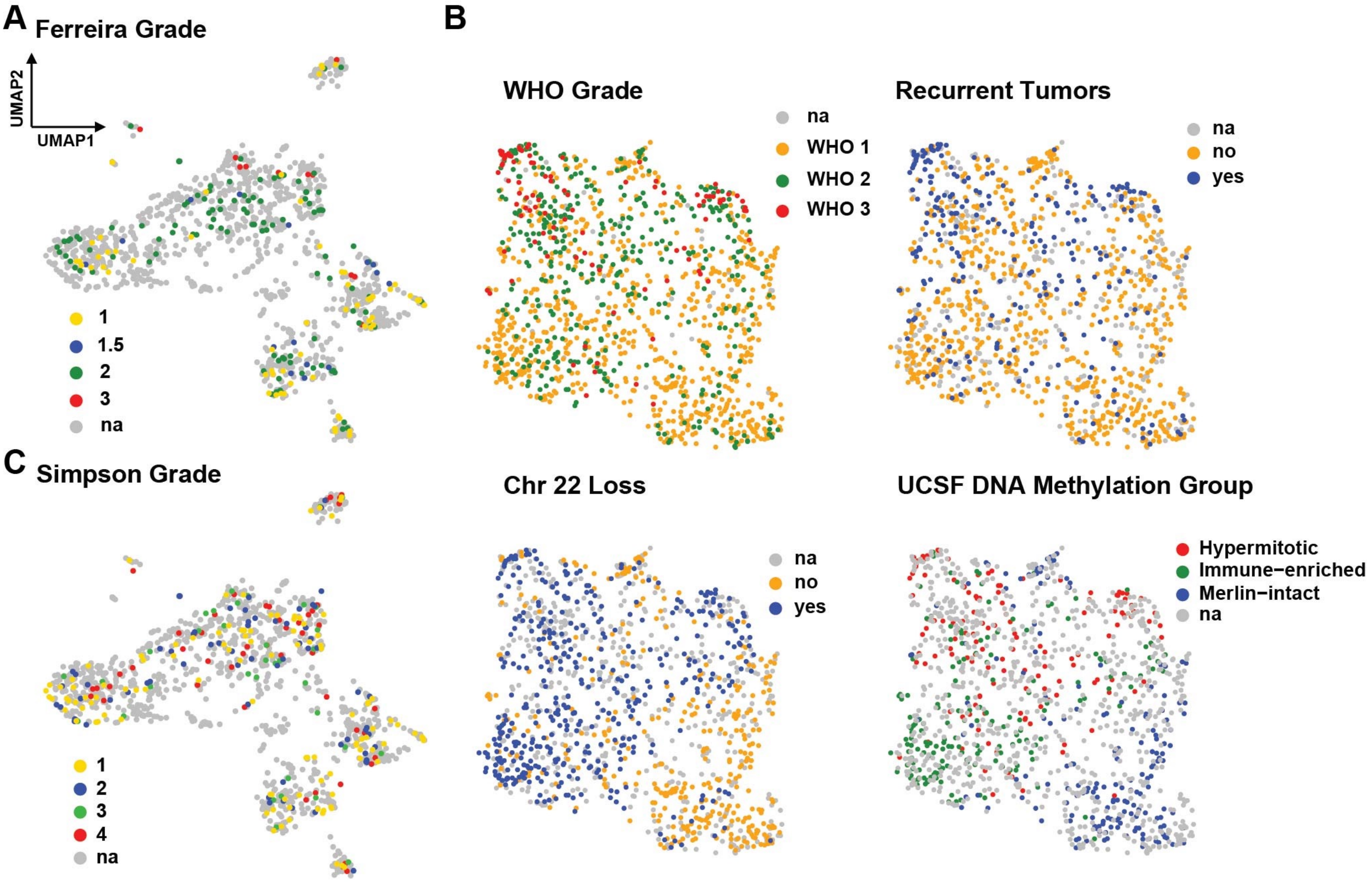

**Figure S3. Related to Figure 3.**

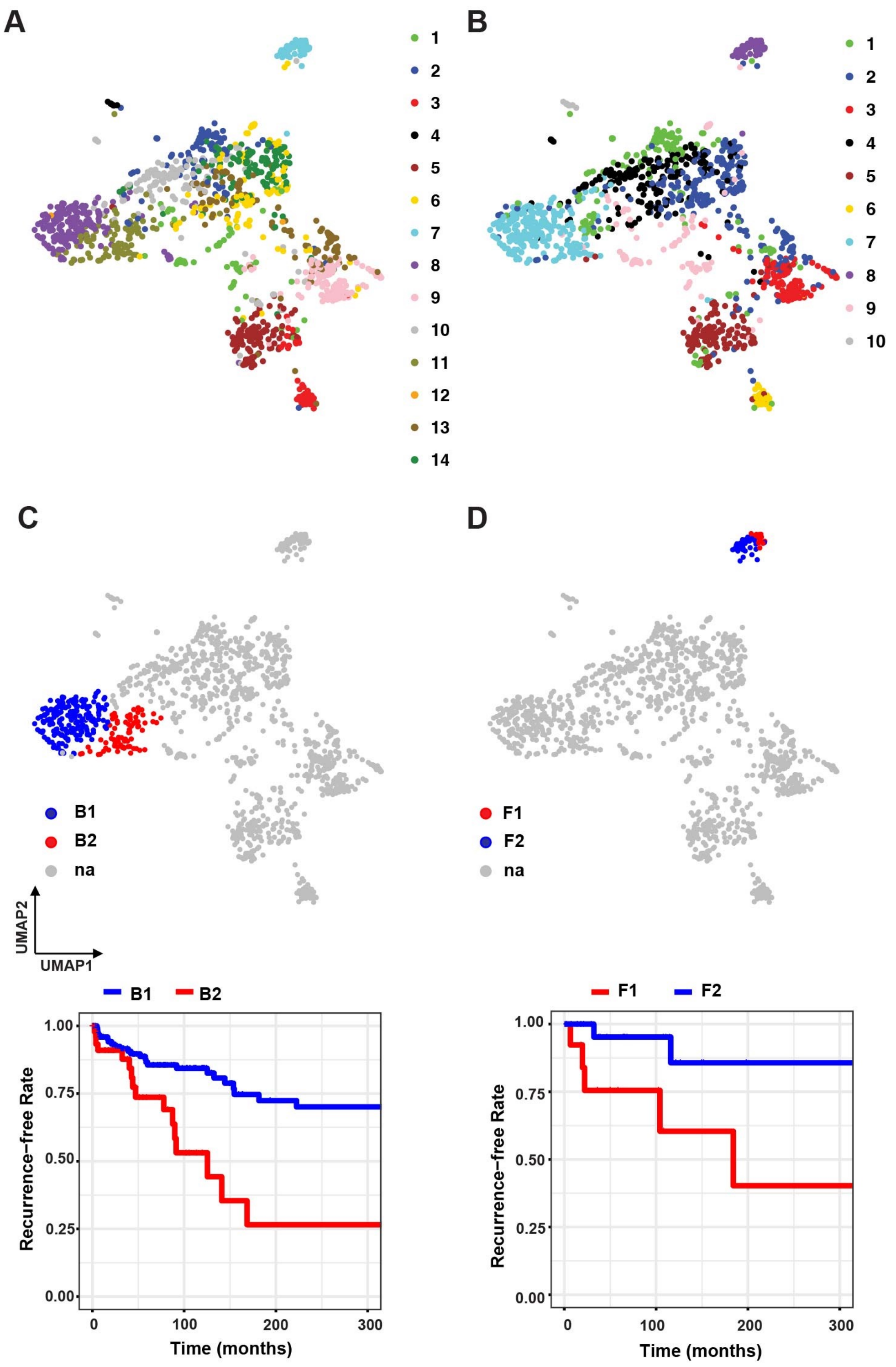

**Figure S4. Related to Figure 4.**

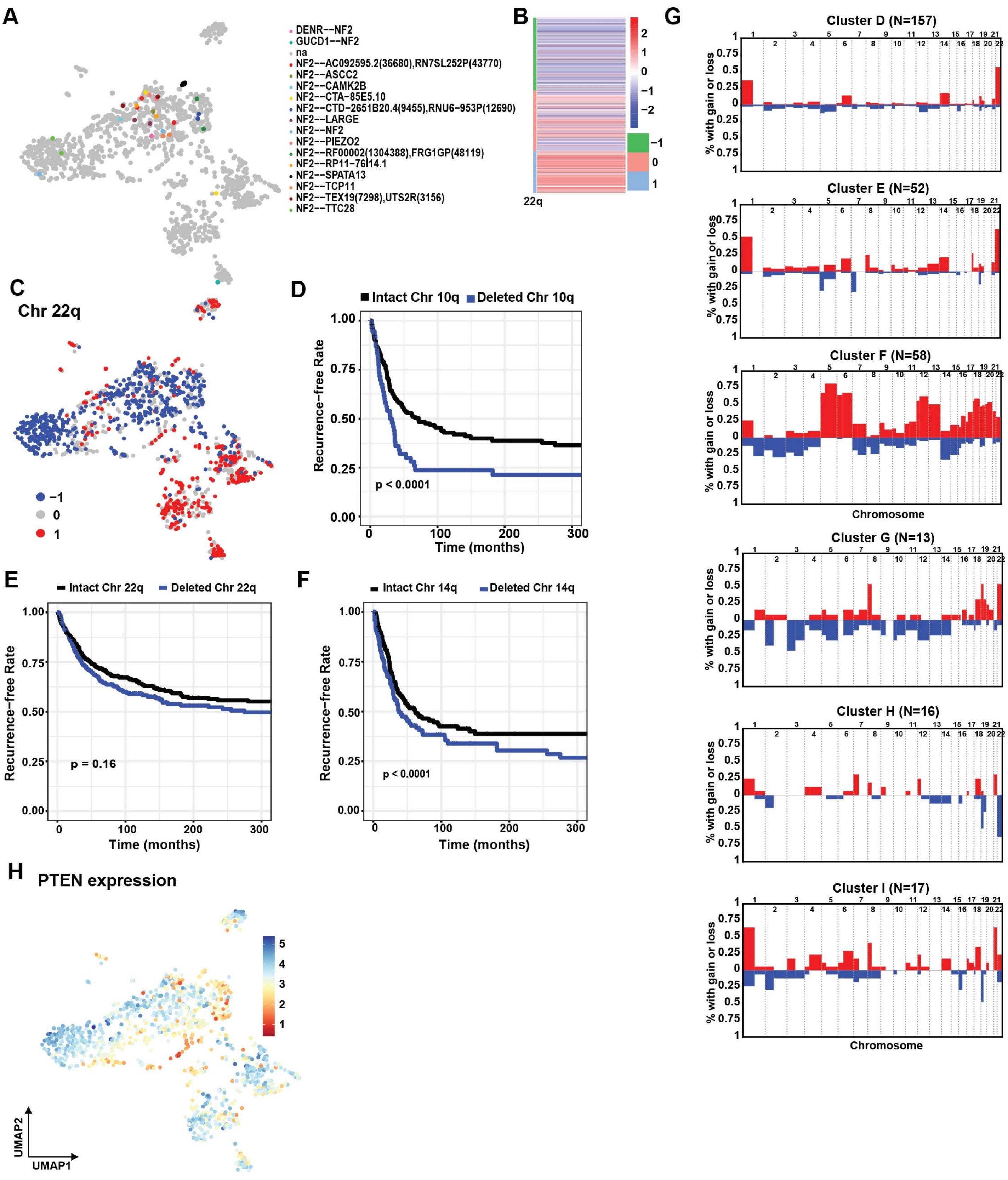

**Figure S4. Related to Figure 4.**

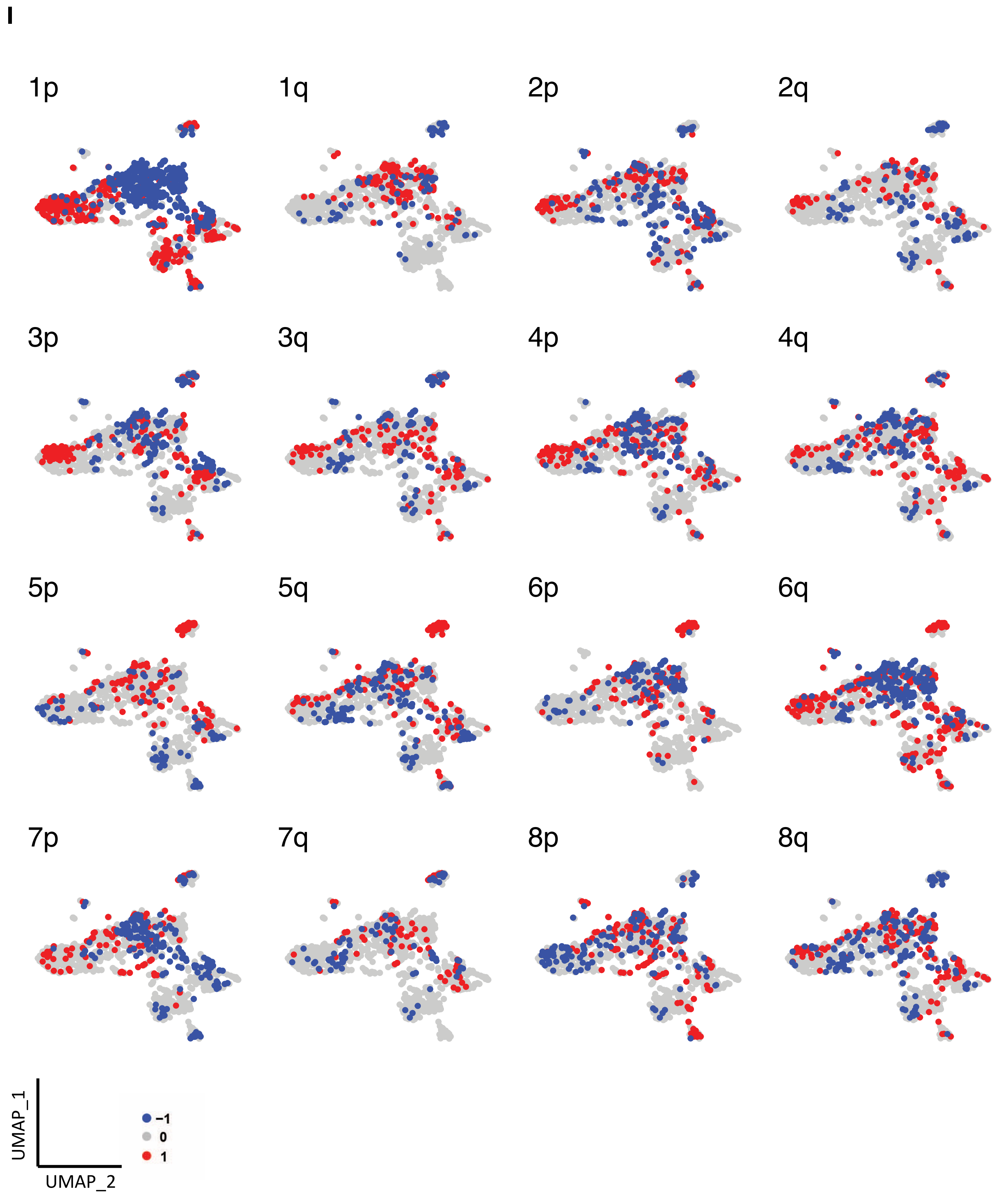

**Figure S4. Related to Figure 4.**

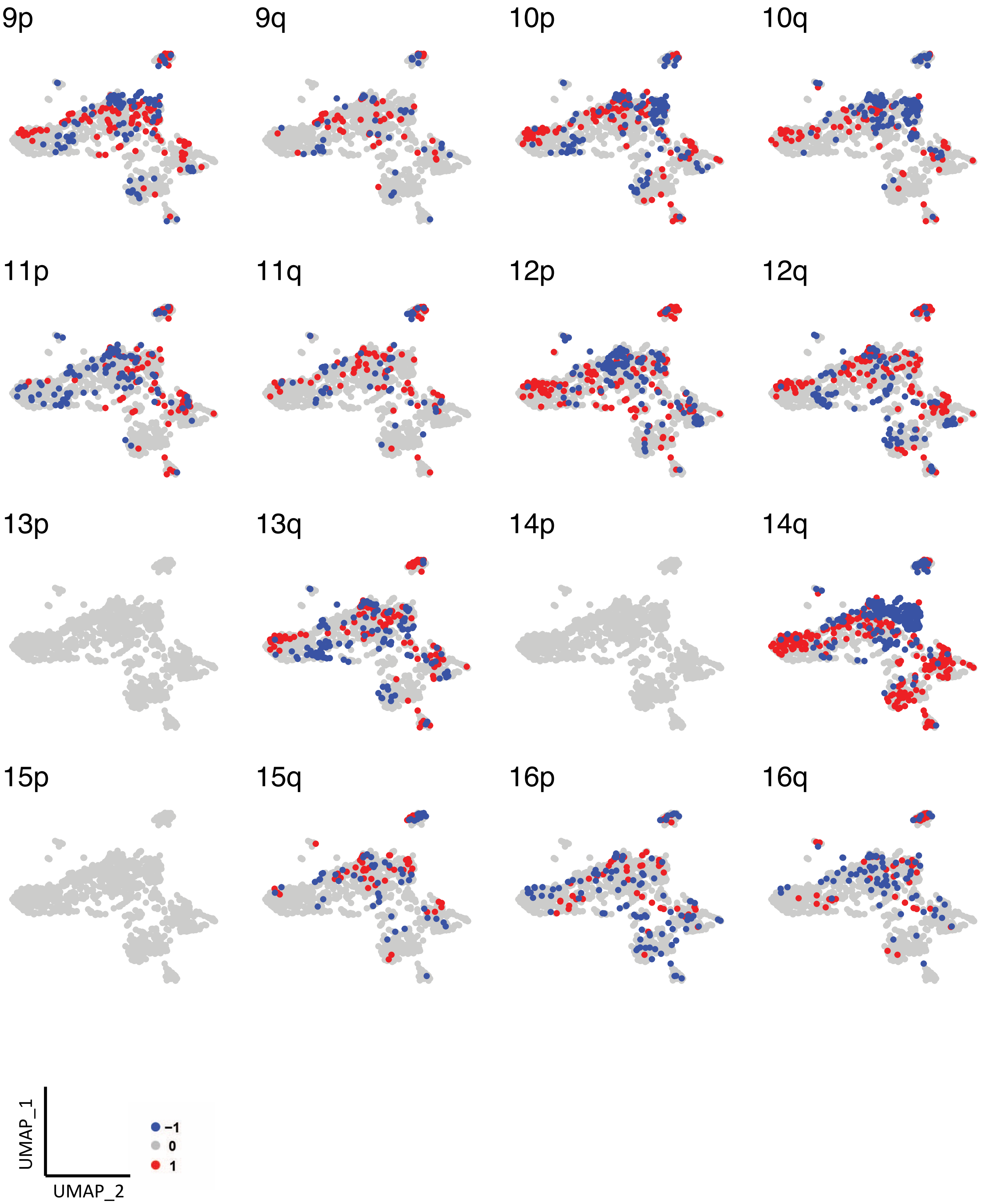

**Figure S4. Related to Figure 4.**

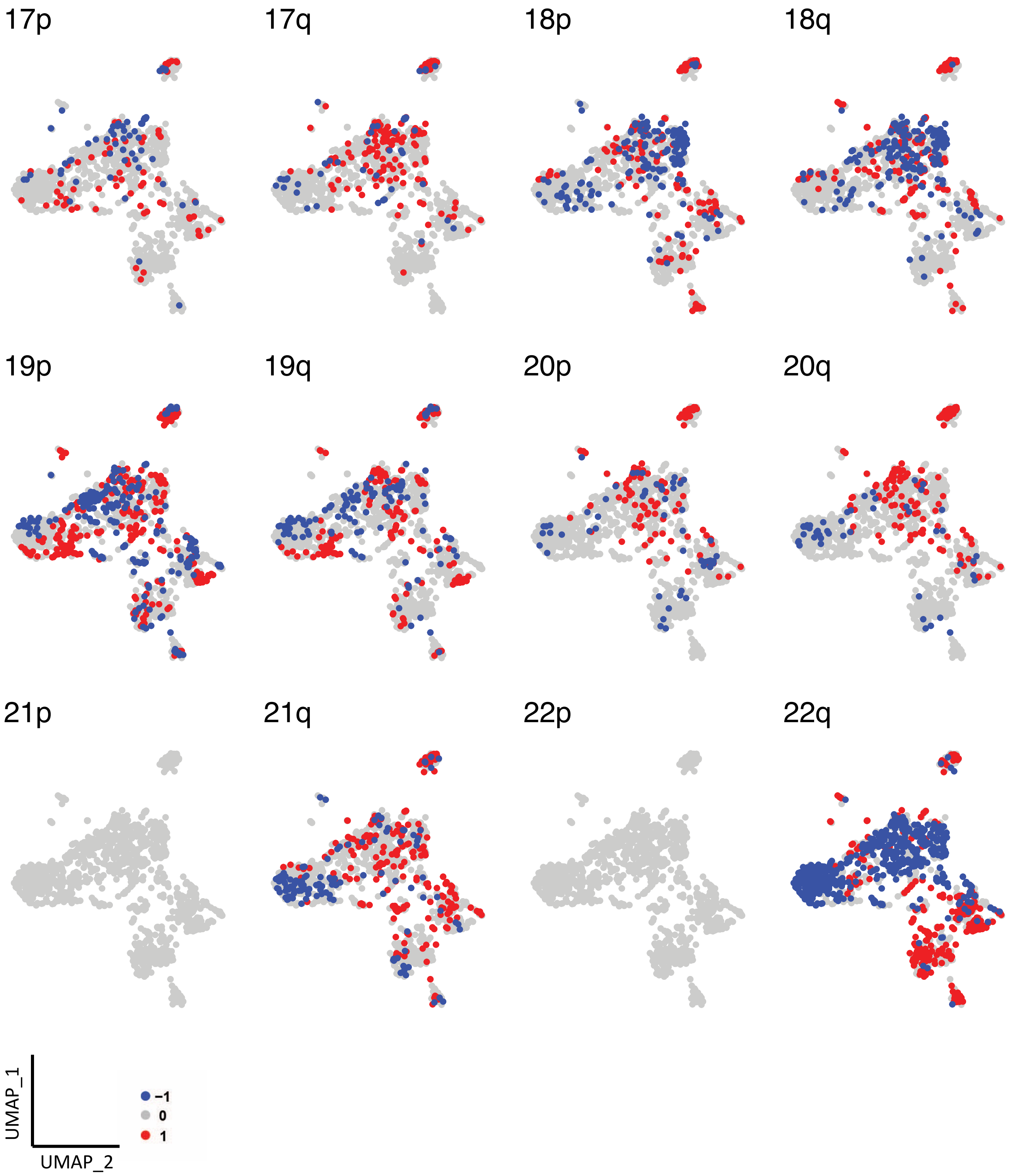

**Figure S5. Related to Figure 5.**

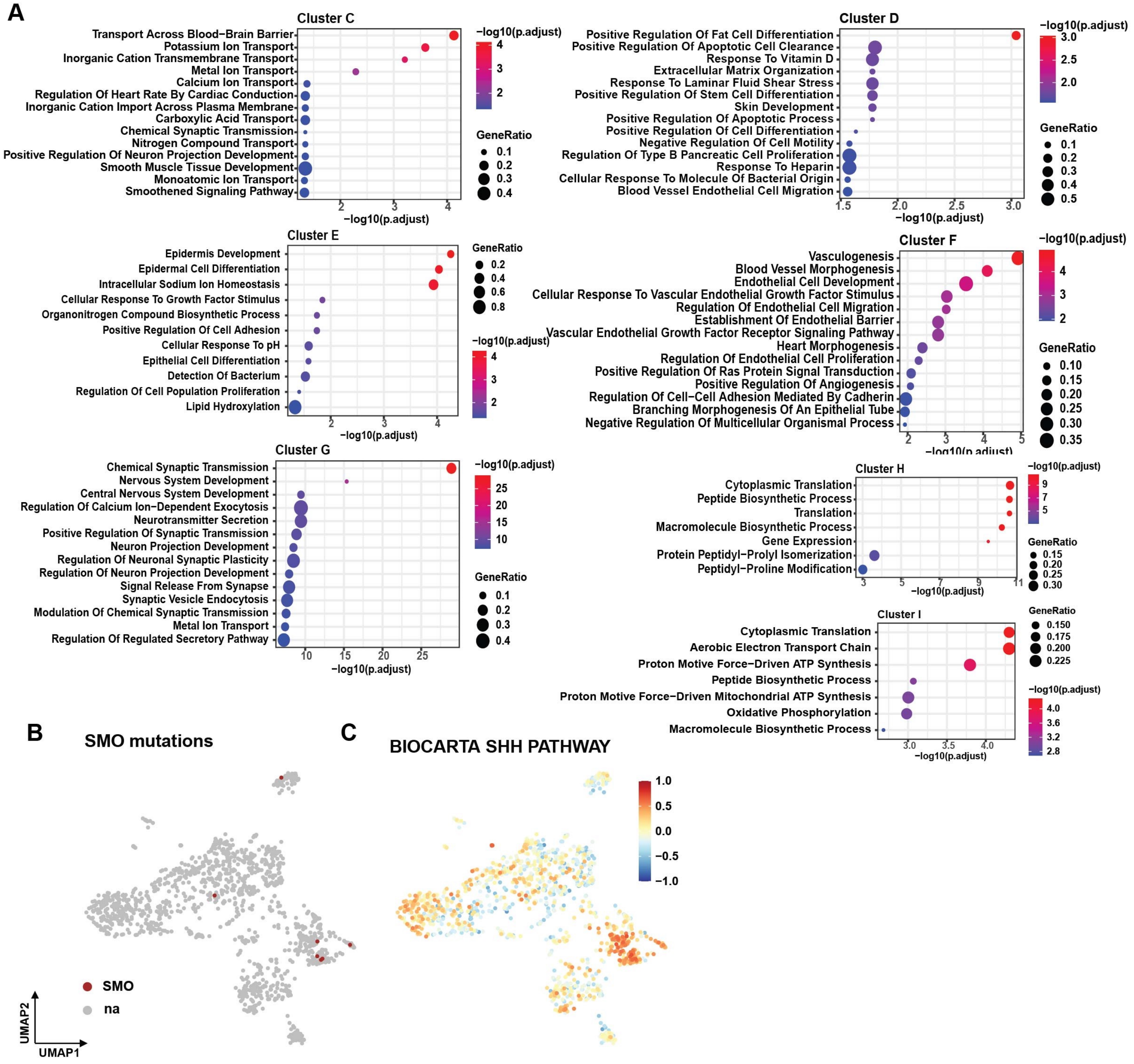

**Figure S5. Related to Figure 5.**
**D**

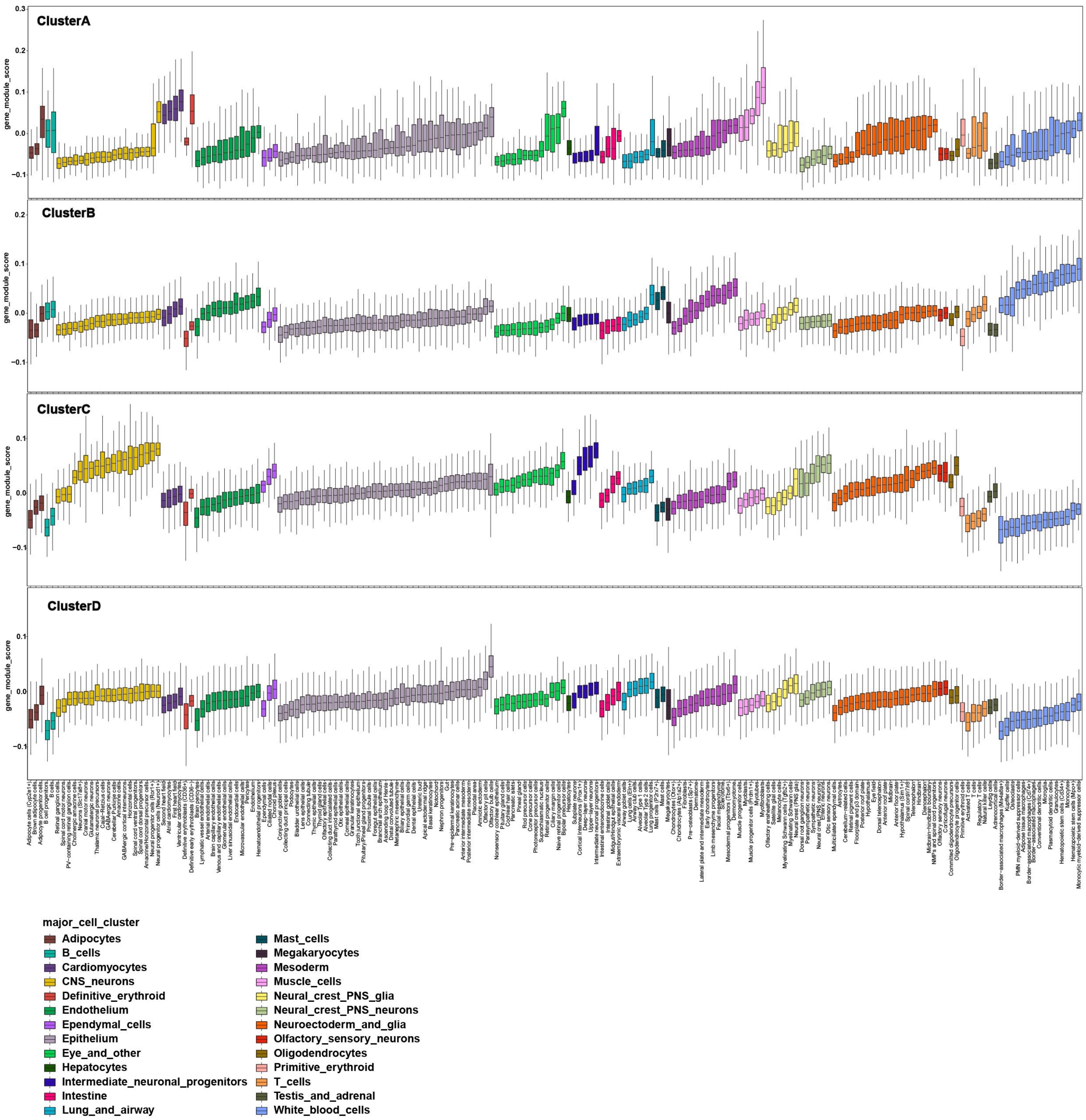

**Figure S5. Related to Figure 5.**

**D**

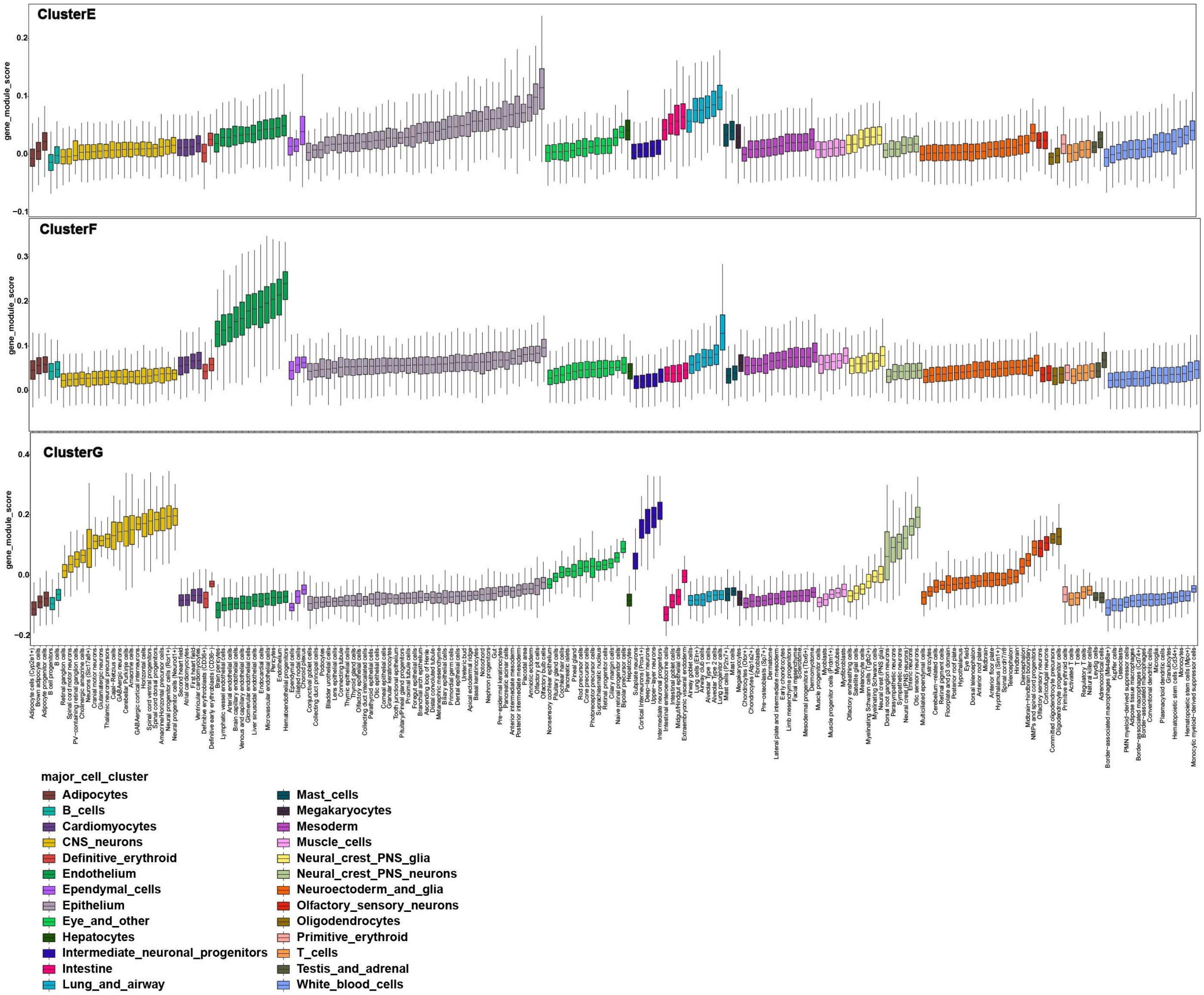

**Figure S7. Related to Figure 7.**

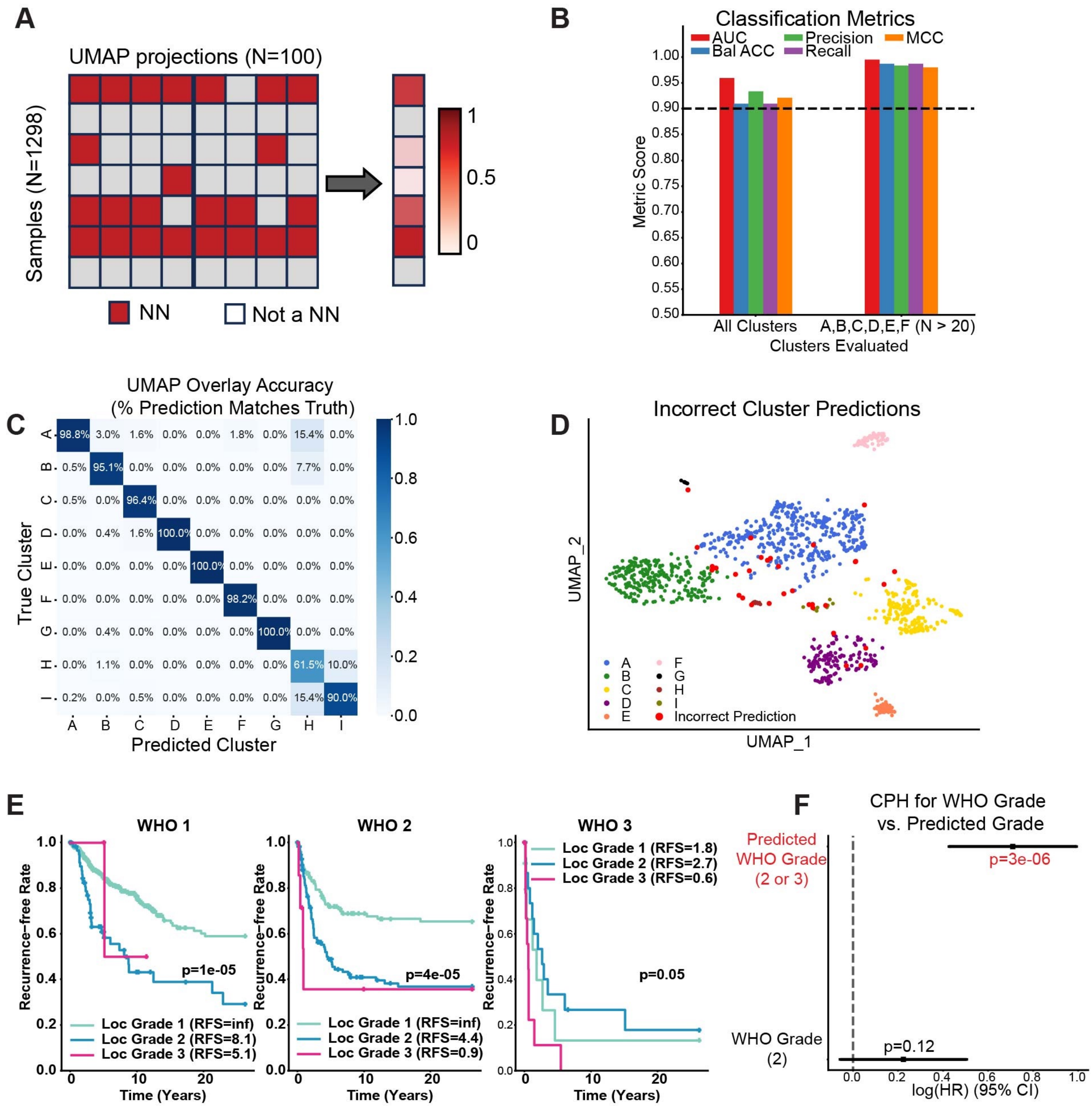

**Table S1. Related to Figure 1. Thirteen bulk RNA-Seq datasets were combined to generate the meningioma UMAP.**

| Dataset | Laboratory | Database Accession Number | Number of samples included in the UMAP | City, Country | Institute |
|---|---|---|---|---|---|
| UW/FHCC | Ferreira/Holland Labs | GSE252291 | 279 | Seattle, USA | University of Washington Fred Hutch Cancer Center |
| Heidelberg | Sahm Lab | -- | 44 | Heidelberg, Germany | University Hospital Heidelberg, Germany |
| UoT | Zadeh Lab | EGAS00001004982 | 123 | Toronto, Canada | University of Toronto, Canada |
| CAVATICA | CAVATICA | CAVATICA | 25 | USA | |
| UCSF(2018) | Raleigh Lab (2018) | GSE101638 | 42 | San Francisco, USA | University of California San Francisco |
| Baylor | Klisch Lab | GSE136661 | 159 | Texas, USA | Baylor College of Medicine |
| UCSD | Rich Lab | GSE139652 | 10 | San Diego, USA | University of California San Diego |
| UCSF(2020) | Raleigh Lab (2020) | GSE151921 | 13 | San Francisco, USA | University of California San Francisco |
| UCSF(2022) | Raleigh Lab (2022) | GSE183656 | 185 | San Francisco, USA | University of California San Francisco |
| Yale | Gunel Lab | GSE85133 | 19 | New Haven, USA | Yale University School of Medicine |
| Palacky | Srovnal Lab | PRJNA705586 | 70 | Czech Republic | Palacky University and University Hospital Olomouc |
| CHOP | CBTN | CBTN | 28 | USA | Children's' Hospital of Philadelphia |
| HKU/UCSF | HKU | GSE212666 | 301 | Hongkong | University of California San Francisco and University of Hong Kong |

**Table S6a. Related to Figure 6. Recurrent tumors in the same patient**

| Patient | Tumor | Age (yrs) | WHO grade | NF2 mutations |
|---|---|---|---|---|
| Patient 1 | 1.1 | 57 | 1 | No |
| | 1.2 | 60 | 1 | No |
| Patient 2 | 2.1 | 33 | 2 | Yes |
| | 2.2 | 42 | 2 | Yes |
| Patient 3 | 3.1 | 47 | 2 | Yes |
| | 3.2 | 61 | 2 | No |
| Patient 4 | 4.1 | 58 | 1 | Yes |
| | 4.2 | 67 | 1 | NA |
| Patient 5 | 5.1 | 45 | 1 | NA |
| | 5.2 | 49 | 1 | NA |
| Patient 6 | 6.1 | 15 | 1 | NA |
| | 6.2 | 20 | 2 | NA |

**Table S6b. Related to Figure 6. Multiple tumors in the same patient**

| Patient | Tumor | Age (yrs) | WHO grade | NF2 mutations |
|---|---|---|---|---|
| Patient 1 | Tumor 1 | 70 | 2 | No |
| | Tumor 2 | 70 | 1 | No |
| Patient 2 | Tumor 1 | 32 | 1 | Yes |
| | Tumor 2 | 34 | 1 | Yes |
| Patient 3 | Tumor 1 | 20 | 1 | Yes |
| | Tumor 2 | 20 | 1 | Yes |
| Patient 4 | Tumor 1 | 18 | 2 | Yes |
| | Tumor 2 | 20 | 2 | Yes |
| | Tumor 3 | 20 | 2 | Yes |

**Table S6c. Related to Figure 6. Progressed tumors in the same patient**

| Patient | Tumor | Age (yrs) | WHO grade |
|---|---|---|---|
| Patient 1 | Progressed 1 | 24 | 2 |
| | Progressed 2 | 25 | 2 |
| Patient 2 | Progressed 1 | 78 | 2 |
| | Progressed 2 | 78 | 2 |
